# Accelerating species loss in southern boreal understories as early warning sign of functional erosion

**DOI:** 10.64898/2026.02.06.704296

**Authors:** Andréa Davrinche, Georg Albert, Elina Kaarlejärvi, Raisa Mäkipää, Tiina Tonteri, Anna-Liisa Laine

## Abstract

Biodiversity is changing rapidly worldwide. Although taxonomic and functional diversity are intrinsically linked, biodiversity change can decouple their trajectories, challenging biodiversity assessments that often focus on a taxonomic perspective. As a result, functional consequences of biodiversity change remain elusive. To better understand how taxonomic and functional diversity change are intertwined, we analytically partition functional diversity change. Specifically, using 38 years of boreal forest understorey data across Finland, we explicitly link species turnover to shifts in community functional composition. We show that, while stable in Mid and Northern regions, recent net loss of fast-growing species in Southern Finland has halted a decades-long increase in functional diversity. This slowdown emerges not from the loss of highly functionally unique species, but rather from the cumulative number of species lost, whose low abundances limit their contribution to functional diversity. Our approach uncovers mechanisms underlying the decoupling of taxonomic and functional diversity, illustrating how functional diversity can persist despite species loss, while also indicating a growing risk of functional erosion. Taken together, these findings underscore the importance and utility of integrating taxonomic and functional dimensions to better anticipate ecological consequences of biodiversity change.

## INTRODUCTION

Anthropogenic environmental change is reshaping biodiversity patterns at an accelerating rate^1^. This change spans multiple dimensions, including high global extinction rates coupled with local turnover in species composition and functional traits^2–4^. Changes in one biodiversity dimension do not necessarily reflect changes in others^5,6^, complicating our ability to evaluate how biodiversity and the functions that it supports are impacted^7^, highlighting the importance of understanding how multiple dimensions of biodiversity are intertwined.

To date, taxonomic diversity, most commonly measured as species richness, is the dimension of diversity most prominently accounted for. However, changes in species richness alone do not always capture the processes underlying biodiversity change^7,8^, or its consequences on the functioning of ecosystems^5,9^. When attempting to link changes in diversity with ecosystem functions, functional diversity (FD), i.e., the diversity of ecological trait values between organisms in a community, has emerged as a key biodiversity metric^5,10^. Capturing the diversity of phenotypes and potential effects on ecosystem functions, functional metrics of diversity enable the generalisation of biodiversity patterns beyond species identity and bring mechanistic insights into the diversity-function spectrum that complement those offered by taxonomic measurements alone^5,7,11–14^.

While taxonomic and functional diversity have been generally assumed to be positively correlated^5,15^, the increasing use of FD alongside taxonomic diversity metrics has brought ample evidence of the context-dependency of this relationship^15,16^. For example, spatially, the link between FD and species richness is known to vary based on geographical scales and associated environmental contexts^17,18^. Further, different facets of FD can respond differently to changes in species richness^13,18^. In particular, because FD accounts for trait similarities between species (i.e., functional redundancy), changes in the diversity of traits can be disconnected from changes in species richness, depending on species uniqueness^5,19,20^.

Temporally, as resident species shift in abundance and species go locally extinct or colonise new habitats, substantial community changes can cancel each other out without being reflected in species richness trends^21,22^. Indeed, globally, local losses and gains of species are accelerating yet mutually offsetting, leading to stable taxonomic richness over time despite important shifts in species identity^23–25^. This species reshuffling can significantly alter community FD, with cascading effects on ecosystem functioning^26^. This discrepancy between taxonomic and functional diversity over time is particularly evident in community dynamics^26–28^. Studies investigating community turnover increasingly consider both taxonomic and functional aspects in parallel^26–29^, with some proposing an integration of both facets of diversity^30,31^. However, a systematic framework translating taxonomic change into its consequences for functional diversity, while retaining essential features of both dimensions of diversity, is still missing.

Quantifying diversity changes and identifying where they emerge from is particularly important for systems facing strong environmental variability and anthropogenic pressure^32–34^. At Northern latitudes, where communities are experiencing fast climatic change^35,36^, boreal forests understories hold a particular ecological interest. They are home to a large share of plant species diversity, and are integral to forest ecosystems^37–39^. In Boreal regions, understorey communities typically form a relatively continuous layer beneath the tree canopy, and play a key role in nutrient cycling, soil processes and habitat provision for fauna^40^. Both climate warming and forest management have accelerated since the last decade, with long-term effects that suggest considering these systems over large time frames to understand their dynamics and predict their future state^41,42^. Hence, boreal forest understories are an ideal system to disentangle how compositional changes affect the community and translates into functional diversity change.

To anticipate and manage biodiversity change, it is crucial to understand the mechanisms driving shifts in natural communities. For this, we link dynamics of community composition to changes in functional diversity by pinpointing the effect of specific species loss, gain and persistence on functional diversity of the community. We base our analysis on the Price equation, which partitions systems’ change into additive components. Initially aimed at quantifying evolutionary changes, the Price equation decomposes changes into their different driving forces^43,44^. Particularly suited for comparing changes between states or time points, the equation has been widely applied across fields, with recent applications in ecology to attribute changes in ecosystem functioning to changes in species diversity^45–50^. Here, we adapt a variation of the equation as developed by Fox & Kerr (2012) to analyse changes in functional diversity over time, by quantifying the functional contribution of each species in a specific community, taking in account their abundance and their functional uniqueness in the community. Hence, beyond quantifying ‘how much diversity’ each species brings to a community, we aim to identify specifically to which extent the effect of changes in community composition on functional diversity is generated from richness changes, or from species’ functional identity and abundance.

Utilising a unique dataset of systematically surveyed boreal vascular understorey vegetation from upland forests across a steep latitudinal gradient of 1 200 km, collected over nearly four decades, we partition the effect of different community composition components on communities’ functional diversity to understand how changes in taxonomic diversity translate into changes in functional diversity (Fig. 1). Specifically, we quantify the relative effect on functional diversity of species turnover (defined here as raw species loss + species gain) and species persistence (i.e., resident species overlapping between two consecutive time points) across time and space. Further, we assess how much of these effects emerge from the number (richness-based effect) or the functional identity (composition-based effect) of the species in the community (Fig. 1), to provide a comprehensive understanding of species’ roles and their temporal changes in upland boreal forest understories.

**Fig 1.**
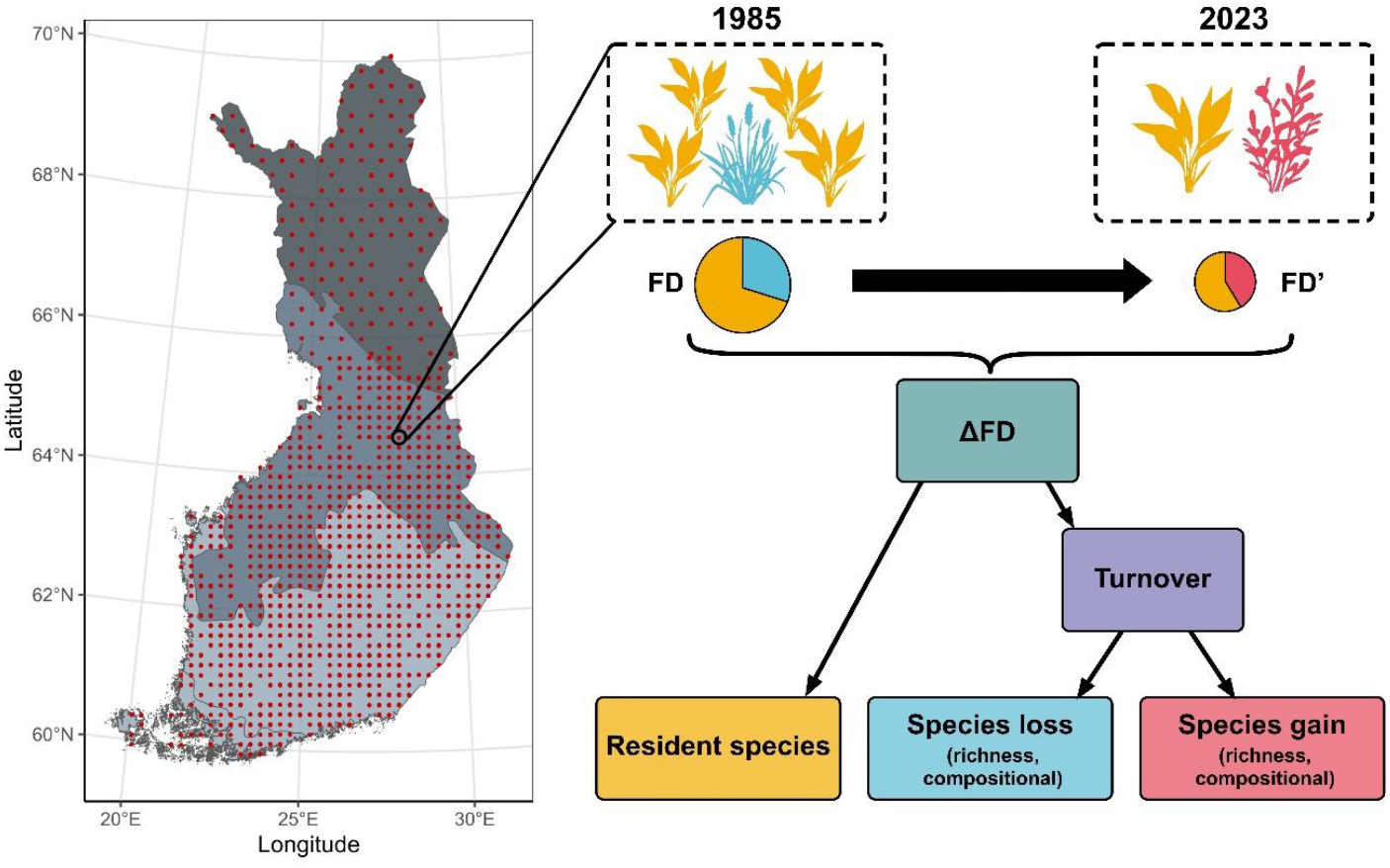
Partitioning of changes in Functional Diversity (ΔFD) across Finnish understorey communities. Between 333 and 1 496 understorey communities of upland forests are compared across consecutive pairwise surveys (1985-1995, 1995-2006, 2006-2023), totalling 1 551 unique communities over 38 years. For each pairwise comparison, changes in functional diversity (ΔFD) of the same community between a baseline (FD) and a comparison (FD’) are partitioned, based on the Price equation (see Methods: Eq. 1). Resulting components describe the contribution to ΔFD of resident species (yellow species) persisting in the community, and of turnover species, with turnover defined here as species loss (blue species; local extinction) and species gain (red species; local colonisation). Species loss and gain are further partitioned into their richness-based and compositional-based components, to distinguish between the influence of species number and species identity in the effect of species turnover on ΔFD.

## RESULTS

### Patterns of taxonomic and functional diversity

Species Richness (SR) and Functional Diversity (FD) of forest understorey communities change with time and depending on the bioclimatic zone (i.e., significant interaction term with *P*<0.001 and *P*=0.001 respectively; Supplementary Table S1). In the South Boreal Zone, SR decreases over the last 17 years with an average loss of 1.310 ± 0.227 species between 2006 and 2023 (*P<*0.001; Supplementary Table S2, Fig. 2A), while SR has otherwise been stable in previous decades in the South, as well as in the rest of the country for the past 38 years.

**Fig 2.**
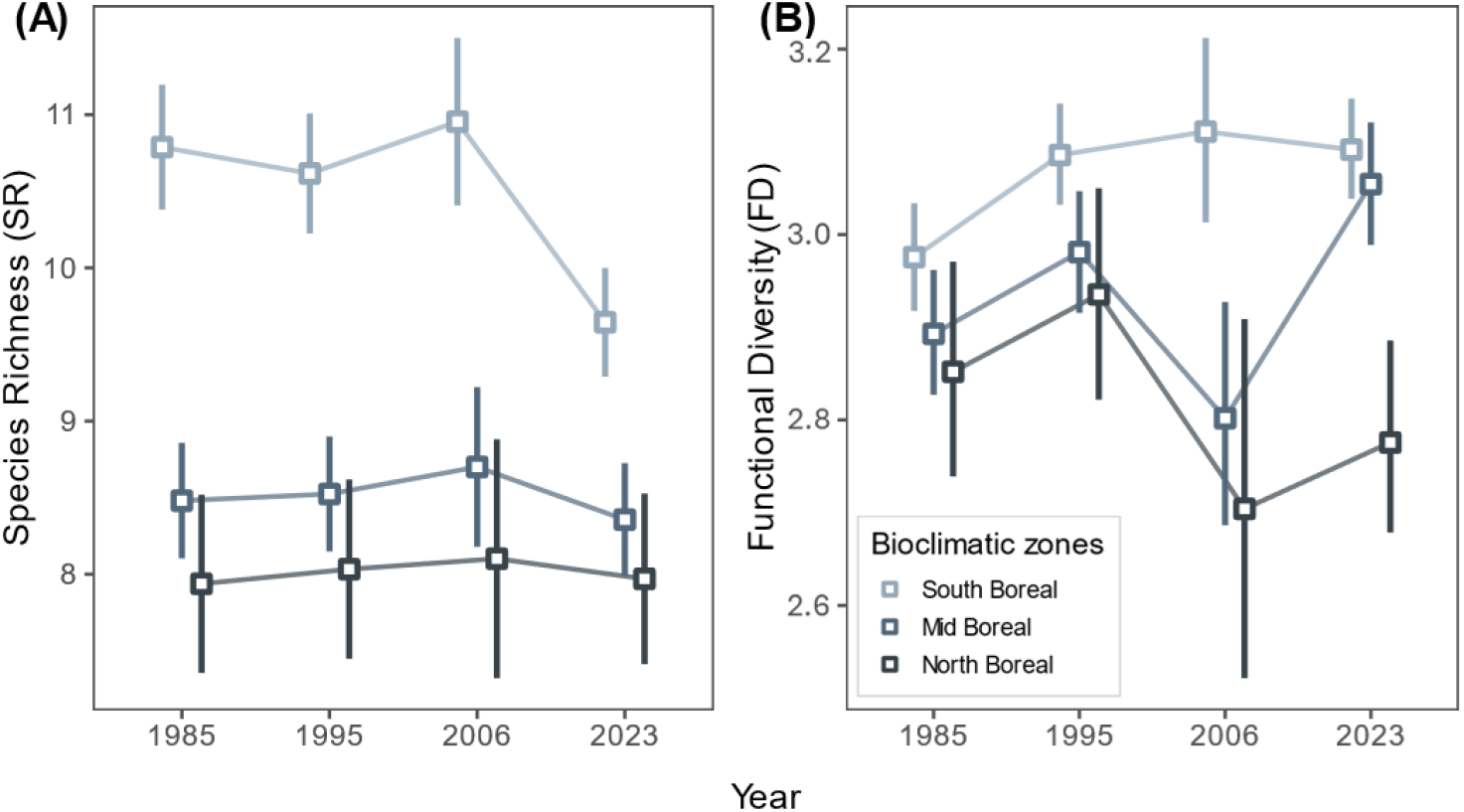
A) Species Richness (SR) and B) Functional Diversity (FD, quantified as Rao’s Q) over time across the South, Mid and North Boreal Zones. Changes in species richness and functional diversity do not show coordinated trends, with species richness loss in the South Boreal Zone not reflected in functional diversity pattern, and stable species richness in the Mid and North Boreal Zones associated with overall increasing and nonsignificant trends of functional diversity respectively. Squares represent the yearly marginal means across sites predicted from linear models (Supplementary Table S1), error bars indicate the 95% confidence interval around the mean. N = 1 589, 1 771, 443 and 1 831 for years 1985, 1995, 2006 and 2023, respectively.

In parallel, FD exhibits different patterns along the latitudinal gradient over time (*F*_*6,4110*_ = 3.74, *P=*0.001; Supplementary Table S1, Fig. 2B). In the South, FD is lower in the first survey year, with significantly higher and more similar values in subsequent years. Meanwhile, FD shows an overall increase in the Mid Boreal Zone, with significantly higher values in 2023 than in 1985. In the North Boreal Zone, changes in FD are not significant across the entire time range (Supplementary Table S2, Fig. 2B).

### Contribution of Resident, Gained and Lost species to changes in FD

Functional diversity, and consequently changes in FD (ΔFD) over time, emerge from three parts of the community: the resident species, persisting through time in the community, the species going locally extinct (species lost) and the species colonising (species gained; Fig. 1). When assessing their relative contribution to ΔFD over time, we find that resident species contribution stays unchanged across the latitudinal gradient but alternates between a positive and negative effect on ΔFD over time (*F*_*2,2358*_ = 9.55, *P<*0.001; Supplementary Table S1, Fig. 3A). On the contrary, species turnover exhibits a stable, close to null contribution in the Mid and North Boreal Zones across time, but shifts into decreasing FD in the South between 2006 and 2023 (*F*_*4,2166*_ = 2.99, *P<*0.05; Supplementary Table S1, Fig. 3A).

**Fig 3.**
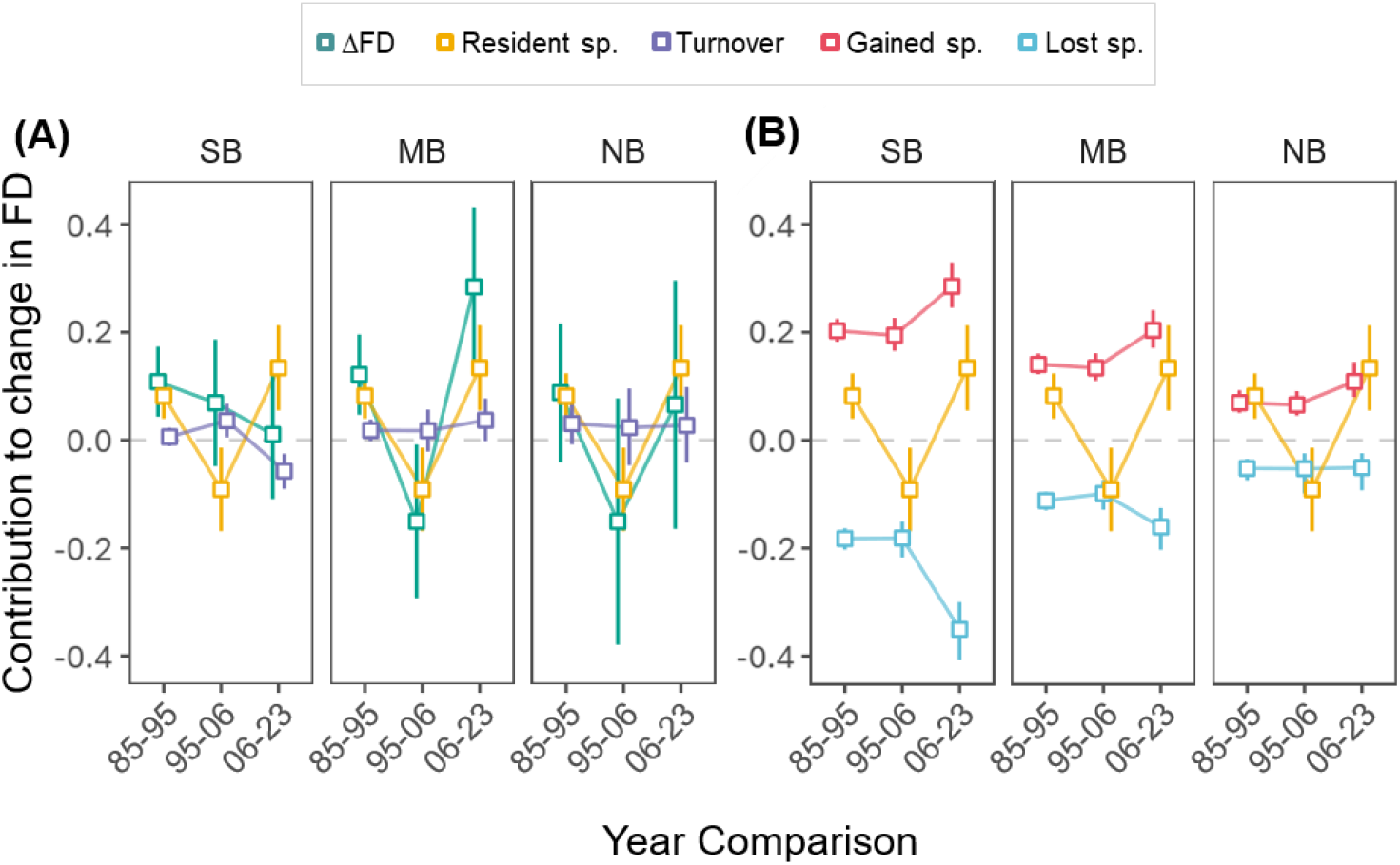
A) Changes in FD (ΔFD) and Contribution to changes in FD of turnover and resident species over time and across South, Mid and North Boreal Zones (SB, MB, NB). Contribution of turnover species decomposes further into B) contribution of species gained and of species lost. ΔFD follows the variable temporal trends of resident species contribution at higher latitudes, while in the South, increase in FD slows down, reflecting the increasing net species loss over time. Squares represent the marginal mean per temporal comparison predicted from a separate linear models per component (Supplementary Table S1), error bars indicate the 95% confidence interval around the mean. The most parsimonious model for each response displays significant effects of the interaction between Year Comparison and Bioclimatic Zone on ΔFD (*P*<0.01), turnover contribution (*P*<0.05) and lost species contribution (*P*<0.05), additive effects of Year Comparison and Bioclimatic Zone on gained species contribution (*P*<0.001; *P*<0.001), and effect of Year Comparison on resident species contribution (*P*<0.001; Supplementary Table S1).

Overall, ΔFD matches closely resident species’ contribution in the Mid and North Boreal Zones. In the South however, although not significant, the rate of increase in FD shows a tendency to decelerate over time, reflecting species turnover’s negative contribution in the last 17 years (Supplementary Table S2, Fig. 3A). When looking separately into the two aspects of turnover, species gain and species loss, we find that the negative contribution of species loss and the positive contribution of species gain to ΔFD gets more pronounced over time. While gain and loss’ contributions mostly compensate each other over time and across Zones, in the South, the contribution of species loss increasingly exceeds that of gain (Fig. 3B).

### Functional contribution of richness and compositional components

To better understand where the effect of gained or lost species on ΔFD originates from, we further decomposed their contribution into what arises from their number (i.e., Richness effect) and what from their identity (i.e., Compositional effect; Fig. 1, Eq. 1, Supplementary Table S8). On the one hand, the richness effect quantifies the effect on ΔFD of losing or gaining an average species, that is, a species that would provide a functional contribution equal to the mean of the community. On the other hand, the compositional effect captures the effect on ΔFD of how different from average the lost or gain species functional contribution is. While in reality neither aspect exists independently from the other, their separation enables us to determine the relative importance of species richness changes (richness effect) compared to changes in species identity (compositional effect).

We find that overall, a change in species number has a greater impact on ΔFD than species identity does, except in the North Boreal Zone where they are of equal importance (Supplementary Fig. S1). While losing a theoretical average species expectedly decreases FD (negative Richness effect of species loss; Supplementary Fig. S1), losing an observed species also tends to have a positive effect on FD when considering only its identity (positive Compositional effect of species loss). This indicates that species lost have a functional contribution lower than the average of the community (same as for species gained, Supplementary Fig. S1). Furthermore, while remaining stable for species gained, species lost functional contribution gets increasingly lower than average over time.

### Effects of functional dissimilarity and species relative abundances

To understand how the species functional profile and presence in the community determines their contribution to change in FD, we further decomposed species-specific functional contribution. Each species functional contribution is defined by its functional dissimilarity to other species in the community and their relative abundances (Eq. 1). We compared these two aspects between lost, gained and resident species to explain differences in contributions to ΔFD.

We find that lost and gained species tend to be functionally less diverse than resident species (Fig. 4A), but do not seem to differ substantially in how distinct they are from the residents (Fig. 4B). Notably, among all species groups, species lost appear most functionally dissimilar from resident species in the Mid Boreal Zone. Species dissimilarity, among group or relative to resident species, remains stable over time, except for species lost whose dissimilarity from resident species marginally increases over time (Fig. 4B, Supplementary Table S5). This comparable dissimilarity of lost and gained species to resident species is also reflected in their trait syndromes, with lost and gained possessing similar trait values, more typical of acquisitive, fast-growing species as opposed to more conservative, slow-growing resident species (Supplementary Fig. S2, Supplementary Tables S6, S7).

**Fig 4.**
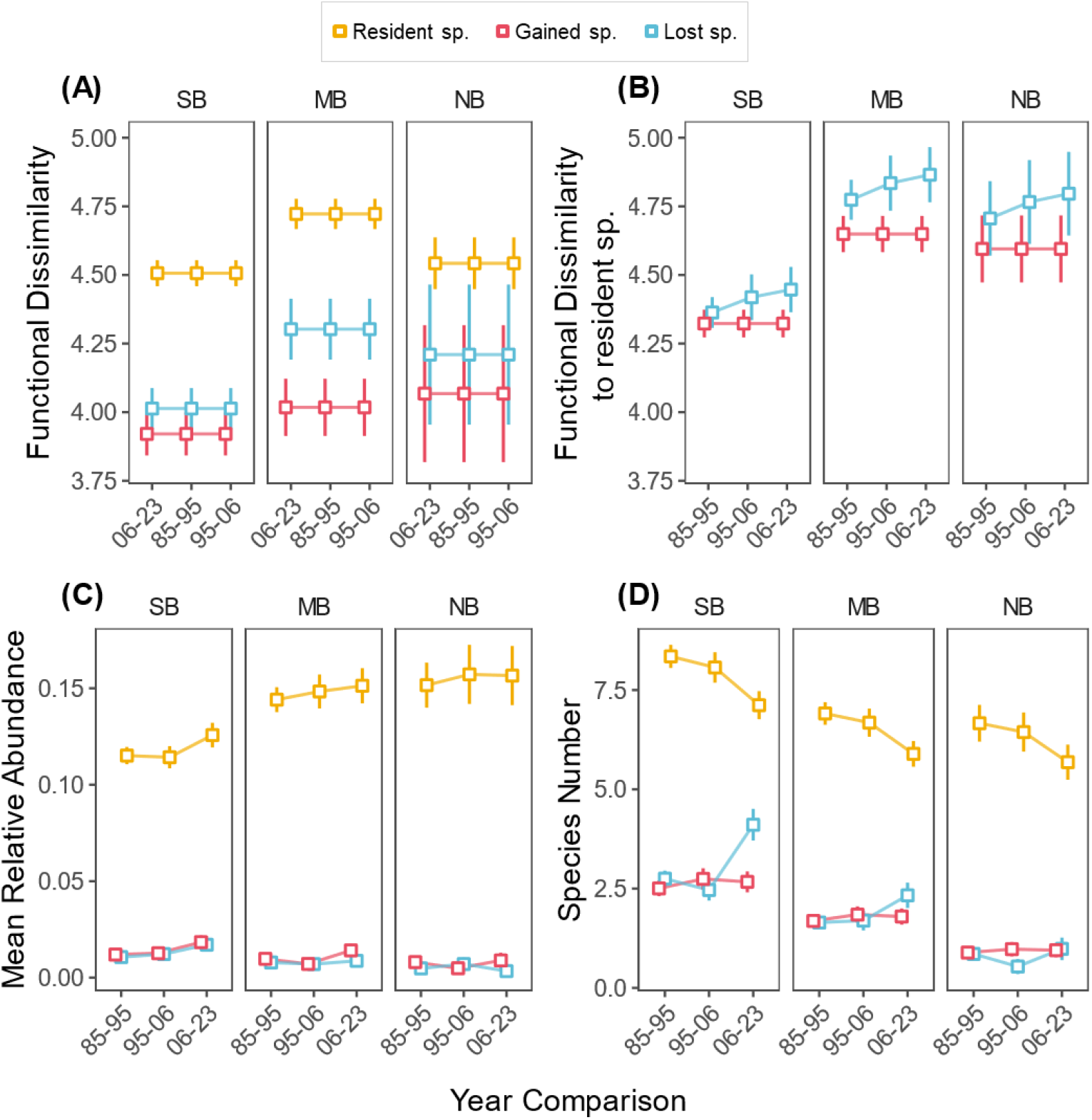
Decomposition of species-specific functional contribution to FD for the three species groups composing the community: resident, lost and gained species. Trends are shown over time and across South, Mid and North Boreal Zones (SB, MB, NB). Functional contribution is decomposed into A) Dissimilarity among resident, lost or gained species, B) Dissimilarity of lost and gained species to resident species, C) species mean relative abundance, and D) species number per group. While resident species are more functionally diverse among themselves, resident, lost and gain are comparably functionally different from each other. They differ more in abundance and number, with resident species increasingly dominating communities over time. Lost and gained species remain few with low abundance, likely resulting in their below-average functional contribution to functional diversity. Squares represent the marginal mean per temporal comparison predicted from a separate linear models per component (Supplementary Table S4), error bars indicate the 95% confidence interval around the mean. The most parsimonious model for each response displays significant effects of A) Bioclimatic Zone on dissimilarity of resident species (*P*<0.001) and lost species (*P*<0.001), B) Bioclimatic Zone on dissimilarity to resident species of lost species (*P*<0.001) and gained species (*P*<0.001), C) additive effects of Year Comparison and Bioclimatic Zone on abundance of resident species (*P*<0.01; *P*<0.001), lost species (*P*<0.05; *P*<0.01) and gained species (*P*<0.001; *P*<0.001), and D) additive effect of Year Comparison and Bioclimatic Zone on number of resident species (*P*<0.001; *P*<0.001), interactive effect of Year Comparison and Bioclimatic Zone on number of lost species (*P*<0.05) and effect of Bioclimatic Zone on number of gained species (*P*<0.001; Supplementary Table S4).

In parallel, the three groups display more marked difference in mean relative abundance and species number, with resident species dominating the communities while lost and gained species reach on average under 3% mean relative abundance and less than 5 species (Fig. 4C & D). Thus, more than their dissimilarity with the rest of the community, the low mean relative abundance of lost and gained species is likely what drives their functional contribution to be lower than the average of the community. Over time, resident species relative abundance increases concurrently to a decrease in species number (Fig. 4C & D, Supplementary Table S5). On the contrary, the number of species gained remains stable while the number of species lost increases over time in all Boreal Zones, with higher losses in the South as reflected in overall Species Richness (Fig. 2A, Fig. 4D, Supplementary Tables S2, S5).

## DISCUSSION

Taxonomic diversity, long central to biodiversity monitoring, does not necessarily mirror changes in functional diversity, which more directly reflects ecosystem processes and resilience^5,10^. Our study of Finland’s boreal understorey communities demonstrates this decoupling clearly: we find that changes in species richness have diverging effects on functional diversity across temporal and spatial scales. Over time, species richness has remained largely stable, except in the in the South Boreal Zone where it declined over the last 17 years, driven by the increasing loss of few, low-abundance, fast-growing species. As a result, functional diversity, which had increased since 1985 due to shifts in the relatively dominant, slow-growing resident species, has begun to level off in recent years in the South. Jointly, our results illustrate that despite a functionally diverse and abundant group of resident species dominating communities’ functional diversity, its temporal stability relies on the balance between turnover and persistence of species. The accentuating loss of increasingly functionally unique species in the South could be an early warning sign of functional diversity erosion.

### Spatio-temporal patterns of taxonomic and functional diversity

Over time, Species Richness (SR) remains stable except for a decline over the last 17 years in the South Boreal Zone. The marked stability in species richness at higher latitudes is consistent with previous reports of stable plant communities’ richness^24,51,52^, including boreal understorey communities^53,54^. However, the recent species loss we observed at lower latitudes contrasts with these findings. Understorey communities in the boreal zone comprise species adapted to harsh environmental conditions, such as low temperatures, abundant snowfall and short growing seasons^42^. Global warming is altering these conditions^35,55^, enabling new species to colonise^51,56^, while potentially driving resident species towards local extinction as their niches shift beyond tolerance limits^57^. As climatic change is generally more pronounced at higher latitudes (e.g., precipitation, snowfall, temperature^35^), these effects would be expected to be strongest in the North. However, we find no evidence for such effects, indicating that the climatic changes were insufficient to show effects in forest understorey communities. This might reflect the decoupling of understorey communities from macroclimate due to forests microclimatic buffering^58^, as well as the time-lag with which species losses is expected to manifest in shrub-dominated northern communities (e.g., ^59,60^). Instead, we found the strongest change in SR in the South, where SR is higher and species loss could result from shifts in biotic interactions. For example, intensified competition, notably between resident and immigrating generalist species^56^, or loss of key interaction partners such as pollinators or mycorrhizal fungi^61,62^ may underlie the observed SR decline. Given stronger human influence in southern Finland, other anthropogenic factors, such as intensifying management practices^63,64^, could also be likely candidates as driver of the observed trends.

In contrast to the stability of SR across most of the period studied, Functional Diversity (FD) exhibited greater temporal variability. At higher latitudes in particular, FD fluctuated over time, whereas in the South the initial rise in FD levelled off. Although the relationship between FD and SR is context-dependent and can vary substantially^16–18^, the number of species strongly constrains FD values, especially in relatively low-diversity communities such as our studied system^5^. However, over the same periods and latitudes, we observed SR remaining largely stable, while to FD varied substantially. This indicates that the shifts in FD must emerge from compositional changes in the community. Hence, considering species dynamics in these communities is a necessary step to understand FD changes driven by species abundances and functional identities, beyond species number.

### Contributions of community dynamics to temporal change in FD

Resident species emerge as the dominant contributors shaping FD trends. With an average positive contribution greater than that of species turnover, they exhibit a variable contribution over the 38 years studied, decreasing, then increasing in more recent years. Resident species’ contribution to changes in FD does not differ across latitudes, suggesting that shared drivers determine its temporal dynamics, such as large-scale macroclimatic trends encompassing the whole region (e.g., amplified climate extremes^42^) or ubiquitous drivers of these community dynamics at the local scale (e.g., forestry management practices, microclimate buffering^65,66^).

In contrast to resident species, the contribution of species turnover to changes in FD is small and remains positive and stable over time and across zones, apart for its decline over the last 17 years in the South Boreal Zone. This small and stable effect emerges from species gain and loss’ respective contribution to changes in FD. Taken independently, species loss and gain contributions are exceeding or comparable to that of resident species. In line with expectations, they also vary with latitude following SR, that is, decreasing in absolute values from South to North^67^. However, because of their opposite effect and comparable values, loss and gain contributions mostly compensate each other. But similar patterns as in the North and Mid Boreal Zone have been reported previously for boreal plant communities, where species richness was found to be generally stable, despite high level of turnover^53,54^, suggesting compensatory losses and gains. In our study, while increasing species gain contribution over time compensates increasing losses in the Mid Boreal Zone, it does not in the South, resulting in a net loss that affects the temporal trend of FD.

Our findings highlight that changes in FD over time are determined by the combination of stabilizing compensatory losses and gain and variable contribution of resident species, all three with equally important independent contributions. Hence, despite the apparent dominating effect of resident species’ contribution to changes in FD, imbalance in species turnover has a direct effect on FD.

### Species-specific functional contribution

To better understand how species contribute to changes in FD based on their role in the community, we further investigated the two aspects that define a species functional contribution: their relative abundance and functional uniqueness. While the first reflects dominance in the community, the latter expresses how functionally unique (i.e., dissimilar) or functionally redundant (i.e., similar) a species is relative to the rest of the community, based on its traits. Greater dissimilarity implies a greater trait space occupied by the community, that is, a greater functional diversity with expected higher ecosystem functioning, resilience and stability over time^10,68^.

On average, resident species represent the most abundant and species-rich group. In addition, they are the most functionally diverse group, compared to functionally homogenous lost and gained species. This high relative abundance combined with comparably high functional uniqueness positions resident species as primary drivers of change in FD. Resident species tend to exhibit more conservative traits, associated with long-lived, slow-growing species, whereas lost and gained species exhibit more acquisitive traits typical of fast-growing, short-lived species. Species with conservative trait values (e.g., high structural values, low leaf nutrient content^69^) intrinsically possess a limited trait adaptability compared to fast growing species^70,71^. Hence, although links between trait syndromes and extinction risks are context-dependent, conservative species might be more prone to extinction in the future, when facing more pronounced change in environmental conditions^72–74^.

In the communities studied, lower adaptability of resident species to change could also result in a shift away from dominant-driven patterns of FD change in the future, toward stronger influence of species turnover. In light of the last 17 years in the South Boreal Zone, we can expect a more prominent role of turnover to lead to an erosion of FD. In line with this, our results also show that over time and across bioclimatic zones, resident species decline in number, suggesting shifts in environmental filtering shaping community composition. Concurrently, lost species exhibit marginally increasing functional uniqueness over time. Hence, species excluded from the communities tend to be those with increasingly unique trait syndromes, reinforcing the idea of an increasingly stringent environmental filter^75,76^, or an accumulation of anthropogenic impacts in their habitat. This is of particular importance in the Mid Boreal Zone, where lost species are the most functionally unique across all groups and zones. While still compensated by gains, species loss increased faster than species gain in the last 17 years, indicating a potential future net loss, and a comparable effect on FD as seen at lower latitudes.

## Conclusion

Integrating functional and taxonomical diversity into a cohesive analytical framework, this study expands the toolset for investigating the consequences of biodiversity change by zooming into communities’ functional composition across space and time. By clarifying the mechanisms that decouple taxonomic and functional diversity, our findings reveal how shifts in community dynamics drive functional diversity change in Finland’s upland boreal forest understories. Importantly, our approach allows the identification of the specific functional identities that contribute to broader temporal patterns of biodiversity change. The observed shifts in community dynamics signal emerging risks for ecosystem future resilience, stability and functioning. Especially in regions facing rapid environmental and land-use change, integrating our approach into existing monitoring schemes would offer a tangible path towards identifying such risks and inform targeted conservation action.

Beyond these immediate implications, our findings point towards several avenues for future work. Our study focused on the overall link between taxonomic and functional diversity change. However, our framework is flexible to the inclusion of alternative functional diversity metrics that would allow to put other aspects into focus, such as communities evenness. Additionally, considering the variability in species’ traits that conditions their adaptability to changing environments presents itself as a promising future step to explore within our framework. Finally, linking trait-based community dynamics to ecosystem functioning will be key to predicting— and potentially mitigating— the consequences of biodiversity loss in a warming world.

## METHODS

### Understorey community data

We use vascular plant species richness, identity and abundance from repeated nationwide vegetation inventories of boreal forest understorey communities conducted by the Natural Resource Institute Finland (LUKE).

The data were collected from a network of 2 043 sites established in forests growing on mineral soil across Finland (Fig. 1). In this network, sites are organised in clusters of three, aligned 600 m apart in northern Finland, and in clusters of four, aligned 400 m apart, in the rest of the country. Clusters are separated by 24 km latitudinally and 32 km longitudinally in the north, and by 16 km in the rest of the country, covering whole Finland. Surveys were carried out in 1985-1986 (referred to as 1985), 1995, 2006 and 2021-2023^77^ (referred to as 2023). In 2006, the survey was conducted as part of a monitoring scheme (BioSoil project, included in the International Co-operative Programme on Assessment and Monitoring of Air Pollution Effects on Forests ^78^) specifically including only one site per cluster. Hence, sample size was reduced for that year, while conserving the extensive spatial coverage of other surveys.

From this pool of sites, all unique 2 043 sites are used to analyse temporal trends of species richness (SR) and functional diversity (FD), across the latitudinal gradient. To be able to analyse site-specific temporal changes in community composition, we compare a same community through time, thus reducing the number of sites used to those common between two consecutive surveys. Hence, we utilize 1 496 sites surveyed both in 1985 and 1995, 441 sites in 1995 and 2006 and 425 sites in 2006 and 2023, for a total of 1 551 unique sites.

For each site, four permanent quadrats of 2 m^2^ were set at 5 m distance from each other. Vascular plant species were identified and their abundance recorded as percentage cover, allowing for species overlap. Cover from the four quadrats was then averaged and converted into relative abundance at site level. The complete dataset included 410 vascular species, excluding tree species saplings. Tree species saplings were excluded from the dataset due to lack of trait data for saplings, which considerably differ from mature trees.

### Functional traits and functional diversity

Nine functional traits representative of growth, dispersal, nutrient acquisition and metabolism were combined from measurements realised in the field (2019-2022) in Finnish boreal forests (following ^79^) and databases sources (TRY, GRoot and NPR, ^80–82^). Measurements carried in the field were prioritized, then completed with trait values from databases. Finally, missing values were imputed using the missForest algorithm ^83^ with a maximum of 10 iterations of 1 000 trees informed by the phylogenetic order of the species (overall NRMSE = 0.396; Supplementary Table S8). Proportion of imputed values was 24.12%, ranging between 2.68% and 67.32% among nine traits (Supplementary Table S8).

### Functional diversity partitioning

Considering each site surveyed in two consecutive time points as a community, we compared changes in community composition between a starting point (i.e., *baseline*, a site at time t) and an end point (i.e., *comparison*, the same site at time t+1) for each single community between 1985 and 1995, 1995 and 2006, and 2006 and 2023. To do this comparison, we partitioned change in the total FD of a community (as Rao’s Q) into richness-based and compositional-based elements of turnover, using a version of the Price equation^46^.

Based on Rao’s Q definition, we used species dissimilarity (calculated as the Euclidean pairwise distance between species) and species relative abundance for each species present in a specific site in a specific year, to calculate the functional contribution of species i, Z_i,_ as follows:

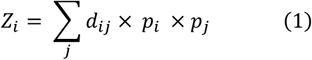

where Z_i_ is the functional contribution of species i, and j the interaction partner of species i, that is, all other species present at the same time as species i in the community; with d_ij_ being the trait-based pairwise dissimilarity between species i and species j, and p_i_ the site- and time-specific relative abundance of each species i in the community. Hence, species functional contribution relates to functional diversity so that *FD* = Σ_*i*_ *Z*.

We calculated the components of the 5-parts Price equation as follow (see Supplementary Table S8 for details):

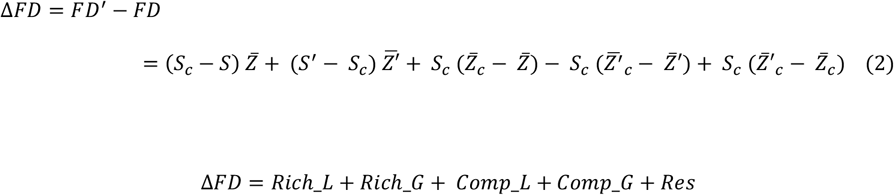

The richness components (Rich_L and Rich_G for loss and gain respectively) capture the impact on the community changes in FD (ΔFD) due to losing or gaining an average species (or *random* species, sensu Fox & Kerr 2012), that is, a species whose functional contribution would be the average of all species in the community. Hence, richness components isolate the effect of species loss or gain on ΔFD due to changes in richness, regardless of the identity of the species. On the contrary, compositional components (Comp_L, Comp_G) capture the impact on the community ΔFD due to losing or gaining a species, with respect to how much their species-specific functional contribution differs from the community’s average. Hence, Comp_L and Comp_G represent the effect of species loss or gain in terms of species identity only, independently from how many species are lost or gained. While existing independently from each other only virtually, richness and compositional components add up together to the impacts of the total loss (Rich_L + Comp_L) and the total gain (Rich_G + Comp_G) of species in a community (Fig. 1).

Because resident species (i.e., species that persist from one time point to the next) do not change in richness or identity between two time points for a same community, Res captures the effect on ΔFD of all changes susceptible to modify the resident species contribution, that is, changes in community composition modifying resident species’ dissimilarity with the remainder of the community, as well as resident species’ abundance.

### Statistical analysis of community change

To analyse change of species richness (SR) and functional diversity (FD) over time, we fitted linear mixed effect models explaining SR or FD by the survey years (Year), the Bioclimatic Zone in which the community is situated, and their interaction. The site identifying code, nested in the site cluster identifying code it belongs to, were used as random intercept to take in account the spatial clustering of sites.

Next, to investigate components of functional diversity change, the following analyses focus on site-specific changes between two consecutive survey years. This includes variables measuring community composition change (resident species, turnover species (i.e., lost and gained), as well as their richness and compositional effects) or changes in community characteristics (functional dissimilarity, relative abundance and number of species of resident, lost or gained species) that contribute to changes in FD (ΔFD). Functional dissimilarity of the different species groups composing the community (i.e., resident, lost and gained species) was analysed both among species of a group and in comparison to resident species. Among species of a group, dissimilarity aimed at evaluating the functional uniqueness of each part of the community independently, by calculating the mean pairwise Euclidean distance between species. For the comparison to resident species, we calculated the mean pairwise Euclidean distance between species lost and resident species, or species gained and resident species, to assess how much uniqueness was being introduced or removed from the community by considering the interaction of turnover species with the rest of the community (i.e., resident species).

Temporal trends of the variables measuring composition change or community characteristics were analysed by fitting linear models with the compared survey years (Year Comparison), the Bioclimatic Zones and their interaction as predictor, with the same random structure as described above (Supplementary Table S1, S3). For turnover, lost and gained species contribution to ΔFD as well as their richness and compositional parts (Rich_L, Rich_G, Comp_L and Comp_G; Eq. 2), zero values represent the absence of lost and/or gained species. As they do not represent an active compositional change in the community, we made the choice not to consider these cases and explicitly remove zero values from these variables. To satisfy to residuals’ normality assumption, SR, lost and gained species dissimilarity among group and compared to resident, as well as the mean relative abundance of all species groups were log-transformed. Lambert W’s transformation was applied to FD, turnover, lost and gained species contribution, and their richness and compositional components. In addition, differences in functional traits values between resident, lost and gained species were tested through linear mixed effect models, with identifying code of site nested in site cluster, further nested in the Bioclimatic Zone as random intercept (Supplementary Table S6).

All models were then simplified stepwise over the fixed and random effects at the threshold of alpha = 0.1, until the most parsimonious model was reached, except for the temporal and latitudinal trends of the dissimilarity within species gained, for which no predictor had a significant effect. For this model, Bioclimatic Zone was kept as unique predictor to facilitate visualisation with other species’ groups (Supplementary Table S5, Fig. 4A). Post-hoc tests were then conducted to determine pairwise differences between temporal comparisons for SR, FD, ΔFD, community composition and community characteristics’ contribution to ΔFD (Supplementary Table S2, S4) variables, as well as for differences in functional traits between species’ groups (Supplementary Table S7), with the Tukey method applied for p-values adjustments.

All analyses were conducted in R version 4.4.2, using *lme4*^84^ for fitting linear mixed effects models and *emmeans*^85^ for estimating marginal means.

## AKNOWLEDGMENTS

The authors acknowledge financial support from Jane and Aatos Erkko Foundation, the Research Council of Finland (grant 362242) and the European Research Council (AdG 101097545 Co-EvoChange) to A.-L.L. We thank the coordinators and surveyors of LUKE for the collection and curation of the vegetation data used in this study.

## DATA AVAILABILITY

All data that support the findings of this study will be made publicly available upon acceptance.

## REFERENCES

1. IPBES. Summary for Policymakers of the Global Assessment Report on Biodiversity and Ecosystem Services. 10.5281/zenodo.3553579 (2019).

2. Keck, F. et al. The global human impact on biodiversity. Nature 641, 395–400 (2025).

3. Pinsky, M. L. et al. Warming and cooling catalyse widespread temporal turnover in biodiversity. Nature 1–5 (2025) doi:10.1038/s41586-024-08456-z.

4. Urban, M. C. Climate change extinctions. Science 386, 1123–1128 (2024).

5. Cadotte, M. W., Carscadden, K. & Mirotchnick, N. Beyond species: functional diversity and the maintenance of ecological processes and services. Journal of Applied Ecology 48, 1079–1087 (2011).

6. Hähn, G. J. A. et al. Global decoupling of functional and phylogenetic diversity in plant communities. Nat Ecol Evol 9, 237–248 (2025).

7. Jarzyna, M. A. & Jetz, W. Detecting the Multiple Facets of Biodiversity. Trends in Ecology & Evolution 31, 527–538 (2016).

8. Vigués Jorba, J. et al. Differential responses of taxonomic, functional and phylogenetic multi-taxa diversity to environmental factors in temperate forest ecosystems. Ecological Indicators 178, 113855 (2025).

9. Le Bagousse-Pinguet, Y. et al. Phylogenetic, functional, and taxonomic richness have both positive and negative effects on ecosystem multifunctionality. Proceedings of the National Academy of Sciences 116, 8419–8424 (2019).

10. Dıaz, S. & Cabido, M. Vive la différence: plant functional diversity matters to ecosystem processes. Trends in Ecology & Evolution 16, 646–655 (2001).

11. Cadotte, M. W., Arnillas, C. A., Livingstone, S. W. & Yasui, S. L. E. Predicting communities from functional traits. Trends in Ecology and Evolution 30, 510–511 (2015).

12. Hooper, D. U. et al. Effects of biodiversity on ecosystem functioning: A consensus of current knowledge. Ecological Monographs 75, 3–35 (2005).

13. Mouchet, M. A., Villéger, S., Mason, N. W. H. & Mouillot, D. Functional diversity measures: an overview of their redundancy and their ability to discriminate community assembly rules. Functional Ecology 24, 867–876 (2010).

14. Petchey, O. L. & Gaston, K. J. Functional diversity: back to basics and looking forward. Ecology Letters 9, 741–758 (2006).

15. Naeem, S. & Wright, J. P. Disentangling biodiversity effects on ecosystem functioning: deriving solutions to a seemingly insurmountable problem. Ecology Letters 6, 567–579 (2003).

16. Morelli, F., Benedetti, Y., Perna, P. & Santolini, R. Associations among taxonomic diversity, functional diversity and evolutionary distinctiveness vary among environments. Ecological Indicators 88, 8–16 (2018).

17. Jarzyna, M. A. & Jetz, W. Taxonomic and functional diversity change is scale dependent. Nat Commun 9, 2565 (2018).

18. Suárez-Castro, A. F., Raymundo, M., Bimler, M. & Mayfield, M. M. Using multi-scale spatially explicit frameworks to understand the relationship between functional diversity and species richness. Ecography 2022, e05844 (2022).

19. Fonseca, C. R. & Ganade, G. Species Functional Redundancy, Random Extinctions and the Stability of Ecosystems. Journal of Ecology 89, 118–125 (2001).

20. Pimiento, C. et al. Selective extinction against redundant species buffers functional diversity. Proceedings of the Royal Society B: Biological Sciences 287, 20201162 (2020).

21. Fletcher Jr., R. J. et al. Beyond Species Richness for Biological Conservation. Conservation Letters 18, e13124 (2025).

22. Larsen, S., Chase, J. M., Durance, I. & Ormerod, S. J. Lifting the veil: richness measurements fail to detect systematic biodiversity change over three decades. Ecology 99, 1316–1326 (2018).

23. Blowes, S. A. et al. The geography of biodiversity change in marine and terrestrial assemblages. Science 366, 339–345 (2019).

24. Dornelas, M. et al. Assemblage Time Series Reveal Biodiversity Change but Not Systematic Loss. Science 344, 296–299 (2014).

25. Dornelas, M. et al. A balance of winners and losers in the Anthropocene. Ecology Letters 22, 847–854 (2019).

26. Kuczynski, L., Bastidas Urrutia, A. M. & Hillebrand, H. Functional diversity loss and taxonomic delays of European freshwater fish and North American breeding birds. Functional Ecology 38, 1726–1738 (2024).

27. Baker, N. J., Pilotto, F., Haubrock, P. J., Beudert, B. & Haase, P. Multidecadal changes in functional diversity lag behind the recovery of taxonomic diversity. Ecology and Evolution 11, 17471–17484 (2021).

28. Robroek, B. J. M. et al. Taxonomic and functional turnover are decoupled in European peat bogs. Nat Commun 8, 1161 (2017).

29. Larson, E. I. et al. A unifying framework for analyzing temporal changes in functional and taxonomic diversity along disturbance gradients. Ecology 102, e03503 (2021).

30. Basile, M. Rare species disproportionally contribute to functional diversity in managed forests. Sci Rep 12, 5897 (2022).

31. Ulrich, W., Zaplata, M. K. & Gotelli, N. J. Reconsidering the Price equation: a new partitioning based on species abundances and trait expression. Oikos 2022, (2022).

32. Daskalova, G. N. et al. Landscape-scale forest loss as a catalyst of population and biodiversity change. Science 368, 1341–1347 (2020).

33. Montràs-Janer, T. et al. Anthropogenic climate and land-use change drive short- and long-term biodiversity shifts across taxa. Nat Ecol Evol 1–13 (2024) doi:10.1038/s41559-024-02326-7.

34. Vellend, M. et al. Plant Biodiversity Change Across Scales During the Anthropocene. Annual Review of Plant Biology 68, 563–586 (2017).

35. Antão, L. H. et al. Climate change reshuffles northern species within their niches. Nat. Clim. Chang. 12, 587–592 (2022).

36. Field, C.B., et al. IPCC, 2014: Climate Change 2014: Impacts, Adaptation, and Vulnerability. (2014).

37. Kumordzi, B. B. et al. Linkage of plant trait space to successional age and species richness in boreal forest understorey vegetation. Journal of Ecology 103, 1610–1620 (2015).

38. Landuyt, D. et al. The functional role of temperate forest understorey vegetation in a changing world. Global Change Biology 25, 3625–3641 (2019).

39. Salemaa, M., Hotanen, J.-P., Oksanen, J., Tonteri, T. & Merilä, P. Broadleaved trees enhance biodiversity of the understorey vegetation in boreal forests. Forest Ecology and Management 546, 121357 (2023).

40. Nilsson, M.-C. & Wardle, D. A. Understory vegetation as a forest ecosystem driver: evidence from the northern Swedish boreal forest. Frontiers in Ecology and the Environment 3, 421–428 (2005).

41. Gauthier, S., Bernier, P., Kuuluvainen, T., Shvidenko, A. Z. & Schepaschenko, D. G. Boreal forest health and global change. Science 349, 819–822 (2015).

42. Venäläinen, A. et al. Climate change induces multiple risks to boreal forests and forestry in Finland: A literature review. Global Change Biology 26, 4178–4196 (2020).

43. Bourrat, P. et al. What is the price of using the Price equation in ecology? Oikos 2023, e10024 (2023).

44. Price, G. R. Selection and Covariance. Nature 227, 520–521 (1970).

45. Fox, J. W. Using the Price equation to partition the effects of biodiversity loss on ecosystem function. Ecology 87, 2687–2696 (2006).

46. Fox, J. W. & Kerr, B. Analyzing the effects of species gain and loss on ecosystem function using the extended Price equation partition. Oikos 121, 290–298 (2012).

47. Winfree, R., W. Fox, J., Williams, N. M., Reilly, J. R. & Cariveau, D. P. Abundance of common species, not species richness, drives delivery of a real-world ecosystem service. Ecology Letters 18, 626–635 (2015).

48. Bannar-Martin, K. H. et al. Integrating community assembly and biodiversity to better understand ecosystem function: the Community Assembly and the Functioning of Ecosystems (CAFE) approach. Ecology Letters 21, 167–180 (2018).

49. Genung, M. A., Fox, J. & Winfree, R. Species loss drives ecosystem function in experiments, but in nature the importance of species loss depends on dominance. Global Ecology and Biogeography 29, 1531–1541 (2020).

50. Ladouceur, E. et al. Linking changes in species composition and biomass in a globally distributed grassland experiment. Ecology Letters 25, 2699–2712 (2022).

51. Thuiller, W., Lavorel, S., Araújo, M. B., Sykes, M. T. & Prentice, I. C. Climate change threats to plant diversity in Europe. Proceedings of the National Academy of Sciences 102, 8245–8250 (2005).

52. Vellend, M. et al. Global meta-analysis reveals no net change in local-scale plant biodiversity over time. Proc Natl Acad Sci U S A 110, 19456–19459 (2013).

53. Happonen, K. et al. Trait-based responses to land use and canopy dynamics modify long-term diversity changes in forest understories. Global Ecol. Biogeogr. 30, 1863–1875 (2021).

54. Kaarlejärvi, E., Salemaa, M., Tonteri, T., Merilä, P. & Laine, A. Temporal biodiversity change following disturbance varies along an environmental gradient. Global Ecol. Biogeogr. 30, 476–489 (2021).

55. Aalto, J. et al. High-resolution analysis of observed thermal growing season variability over northern Europe. Clim Dyn 58, 1477–1493 (2022).

56. Sommer, J. H. et al. Projected impacts of climate change on regional capacities for global plant species richness. Proceedings of the Royal Society B: Biological Sciences 277, 2271–2280 (2010).

57. Villén-Peréz, S., Heikkinen, J., Salemaa, M. & Mäkipää, R. Global warming will affect the maximum potential abundance of boreal plant species. Ecography 43, 801–811 (2020).

58. De Frenne, P. et al. Microclimate moderates plant responses to macroclimate warming. Proceedings of the National Academy of Sciences 110, 18561–18565 (2013).

59. González-Varo, J. P., Albaladejo, R. G., Aizen, M. A., Arroyo, J. & Aparicio, A. Extinction debt of a common shrub in a fragmented landscape. Journal of Applied Ecology 52, 580–589 (2015).

60. Vellend, M. et al. Extinction Debt of Forest Plants Persists for More Than a Century Following Habitat Fragmentation. Ecology 87, 542–548 (2006).

61. Laine, A.-L. & Tylianakis, J. M. The coevolutionary consequences of biodiversity change. Trends in Ecology & Evolution 39, 745–756 (2024).

62. Sandor, M. E., Elphick, C. S. & Tingley, M. W. Extinction of biotic interactions due to habitat loss could accelerate the current biodiversity crisis. Ecological Applications 32, e2608 (2022).

63. Kuuluvainen, T. & Siitonen, J. Fennoscandian boreal forests as complex adaptive systems: Properties, management challenges and opportunities. in Managing Forests as Complex Adaptive Systems (Routledge, 2013).

64. Määttänen, A.-M., Virkkala, R., Leikola, N. & Heikkinen, R. K. Increasing loss of mature boreal forests around protected areas with red-listed forest species. Ecol Process 11, 17 (2022).

65. De Frenne, P. et al. Forest microclimates and climate change: Importance, drivers and future research agenda. Global Change Biology 1–19 (2021) doi:10.1111/gcb.15569.

66. Nirhamo, A., Aakala, T. & Kouki, J. Forest biodiversity in boreal Europe: Species richness and turnover among old-growth forests, managed forests and clearcut sites. Biological Conservation 306, 111147 (2025).

67. Hillebrand, H., Soininen, J. & Snoeijs, P. Warming leads to higher species turnover in a coastal ecosystem. Global Change Biology 16, 1181–1193 (2010).

68. de Bello, F. et al. Functional trait effects on ecosystem stability: assembling the jigsaw puzzle. Trends in Ecology & Evolution 36, 822–836 (2021).

69. Reich, P. B. The world-wide ‘fast-slow’ plant economics spectrum: A traits manifesto. Journal of Ecology 102, 275–301 (2014).

70. Böhnke, M. & Bruelheide, H. How do evergreen and deciduous species respond to shade?— Tolerance and plasticity of subtropical tree and shrub species of South-East China. Environmental and Experimental Botany 87, 179–190 (2013).

71. Gorné, L. D. et al. The acquisitive–conservative axis of leaf trait variation emerges even in homogeneous environments. Ann Bot 129, 709–722 (2020).

72. Chichorro, F. et al. Trait-based prediction of extinction risk across terrestrial taxa. Biological Conservation 274, 109738 (2022).

73. Maracahipes-Santos, L. et al. Intraspecific trait variability facilitates tree species persistence along riparian forest edges in Southern Amazonia. Sci Rep 13, 12454 (2023).

74. Purvis, A., Jones, K. E. & Mace, G. M. Extinction. BioEssays 22, 1123–1133 (2000).

75. Gaüzère, P. et al. The functional trait distinctiveness of plant species is scale dependent. Ecography 2023, e06504 (2023).

76. Padullés Cubino, J. Environmental drivers of taxonomic and functional turnover of tree assemblages in Europe. Oikos 2023, e09579 (2023).

77. Tonteri, T. et al. Operation Bilberry: Inventory of the forest vegetation on permanent plots in Finland in 2021 - 2023. Natural Resources Institute Finland. fairdata.fi (2026).

78. Lorenz, M. International Co-operative Programme on Assessment and Monitoring of Air Pollution Effects on Forests-ICP Forests-. Water Air Soil Pollut 85, 1221–1226 (1995).

79. Pérez-Harguindeguy, N. et al. New handbook for standardised measurement of plant functional traits worldwide. Australian Journal of Botany 61, 167–234 (2013).

80. Guerrero-Ramírez, N. R. et al. Global root traits (GRooT) database. Global Ecology and Biogeography 30, 25–37 (2021).

81. Kattge, J. et al. TRY plant trait database – enhanced coverage and open access. Global Change Biology 26, 119–188 (2020).

82. Tian, D. et al. A global database of paired leaf nitrogen and phosphorus concentrations of terrestrial plants. Ecology 100, e02812 (2019).

83. Stekhoven, D. J. & Bühlmann, P. MissForest—non-parametric missing value imputation for mixed-type data. Bioinformatics 28, 112–118 (2012).

84. Bates, D., Maechler, M., Bolker, B. & Walker, S. Fitting Linear Mixed-Effects Models Using lme4. Journal of Statistical Software 1, 1–48 (2015).

85. Lenth, R. & Piaskowski, J. emmeans: Estimated Marginal Means, aka Least-Squares Means. (2025).

