## Supplementary Materials for "Accelerating species loss in southern boreal understories as early warning sign of functional erosion"

### Figures

Fig. S1. Five-parts Price equation components: resident species, and lost and gained species' Richness and Compositional contributions to FD over time.

Fig. S2. Average functional traits values of community's species groups (resident, lost and gained species).

### Tables

Table S1. Model results for Species Richness (SR) and Functional Diversity (FD) over time and Bioclimatic Zones, and Changes in FD (ΔFD), Turnover and Resident species' contribution to ΔFD (Res.) over consecutive survey years (Year Comparison) and Bioclimatic Zones (Fig. 2, 3 & Fig. S1).

Table S2. Post-hoc test results for models of Species Richness (SR), Functional Diversity (FD), Changes in FD (ΔFD), Turnover species' contribution to ΔFD, and Resident species' contribution to ΔFD (Res.) (Fig. 2, 3 & Fig. S1).

Table S3. Five-parts Price equation components description.

Table S4. Model results for species groups' (resident, lost and gained species) Dissimilarity, Relative Abundance and Number over consecutive survey years (Year Comparison) and Bioclimatic Zones (Fig. 4).

Table S5. Post-hoc test results for models of species groups' (resident, lost and gained species) Dissimilarity, Relative Abundance and Number (Fig. 4).

Table S6. Model results for functional traits differences between species groups (resident, lost and gained species; Fig. S2).

Table S7. Post-hoc test results for models of functional traits differences between species groups (resident, lost and gained species; Fig. S2).

Table S8. Functional traits range and imputation errors.

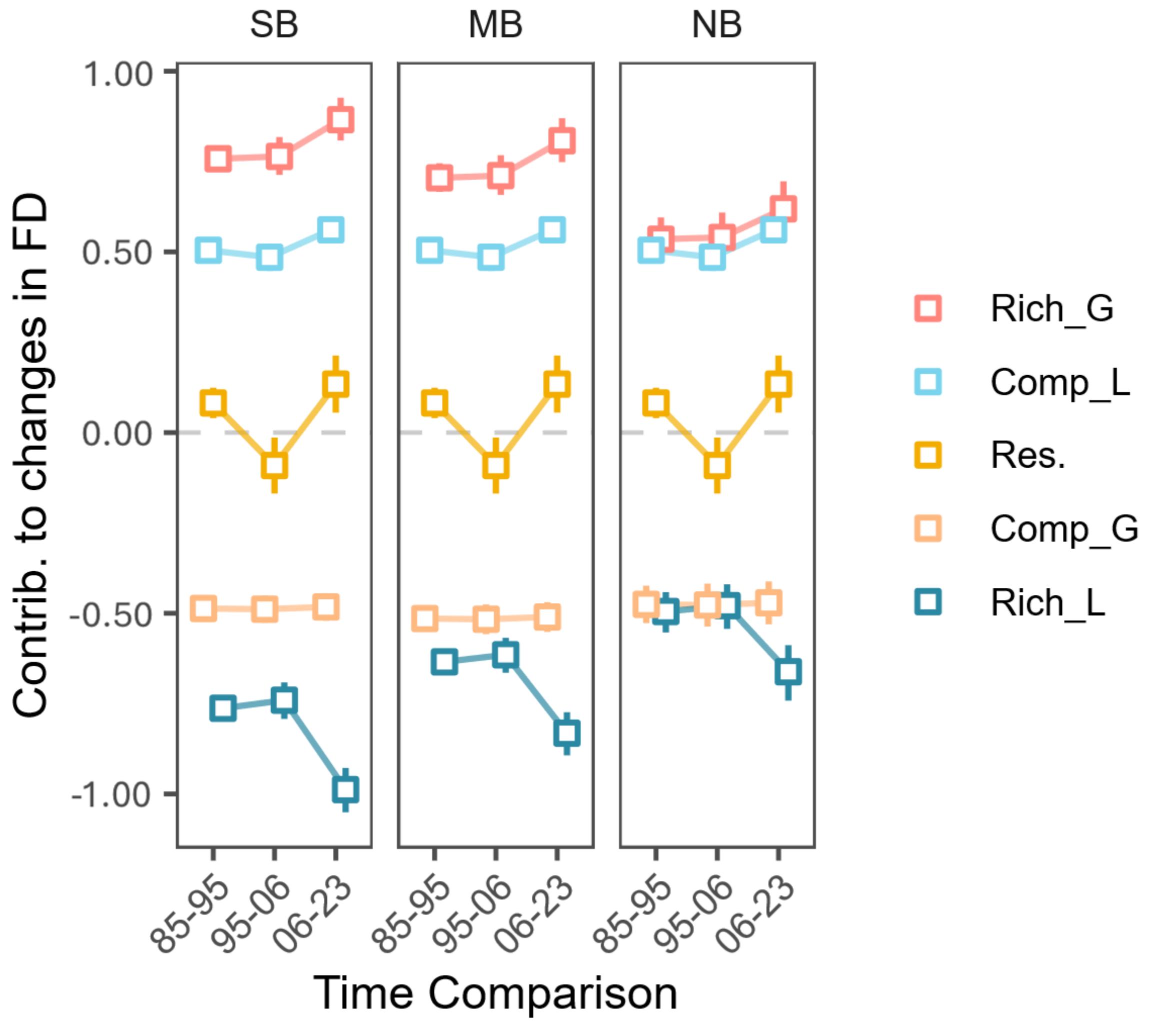


**Fig. S1. Five-parts Price equation components: resident species, and lost and gained species' Richness and Compositional contributions to FD over time (Res., Rich_L, Rich_G, Comp_L, Comp_G; see Eq. 1, Supplementary Table S3).** Magnitude of richness effects exceeds that of compositional effects, and increases over time. Compositional effects reveal lost and gained species contribution to be below average compared to the rest of the community. Squares represent the yearly marginal means across sites predicted from linear models (Supplementary Table S1), error bars indicate the 95% confidence interval around the mean. The most parsimonious model for each response displays significant effects of Year Comparison and Bioclimatic Zone on the contribution to ΔFD of gained species richness effect (*P*=0.001; P<0.001) and lost species richness effect (*P*<0.001; *P*<0.001), and a significant effect of Year Comparison on lost species compositional effect (*P*<0.01), and on resident species contribution (*P*<0.001; Supplementary Table S1).

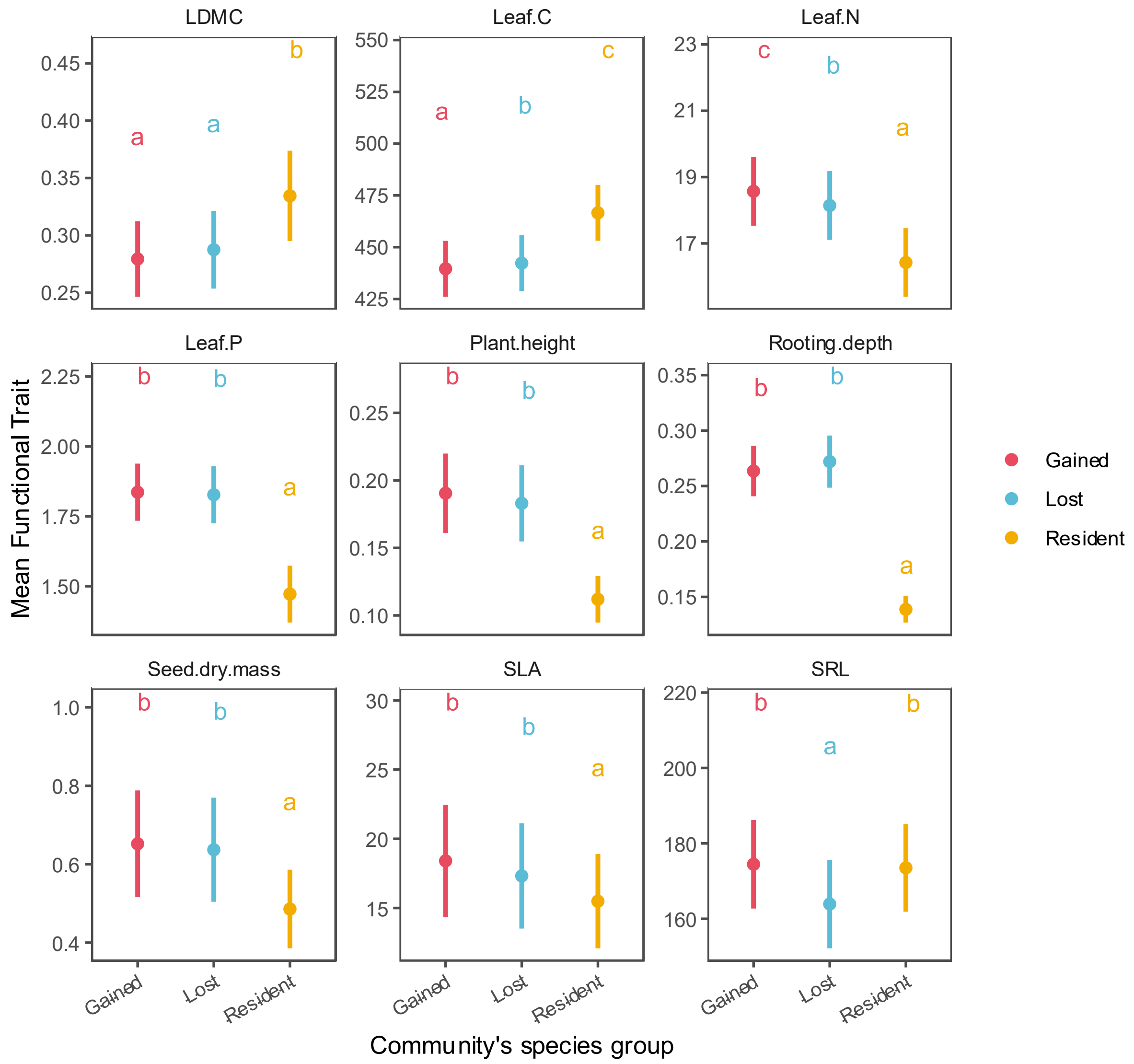


**Fig. S2. Average functional traits values of community's species groups (resident, lost and gained species).** While resident species possess traits more typical for conservative, slow-growing species, lost and gained species differ little in their fast-growing, acquisitive functional profile. Circles represent the marginal means over time and across sites predicted from linear models for each trait (Table S6), error bars indicate the 95% confidence interval around the mean and letters represents significance of differences between means at alpha = 0.05 (Table S7). See Table S6 for trait abbreviations.

**Table S1**. Model results for Species Richness (SR) and Functional Diversity (FD) over time and Bioclimatic Zones, and Changes in FD (ΔFD), Turnover and Resident species' contribution to ΔFD (Res.) over consecutive survey years (Year Comparison) and Bioclimatic Zones (Fig. 2, 3 & S1).

| Response | Predictor | Sum Sq. | Mean Sq. | Num.DF | Den.DF | F-value | P-value |
|---|---|---|---|---|---|---|---|
| Species Richness (SR) | | | | | | | |
| | Year | 1.592 | 0.531 | 3 | 3723.064 | 7.858 | < 0.001 |
| | Bioclimatic Zone | 5.468 | 2.734 | 2 | 1022.196 | 40.493 | < 0.001 |
| | Year:Bioclimatic Zone | 2.422 | 0.404 | 6 | 3722.785 | 5.978 | < 0.001 |
| Functional Diversity (FD) | | | | | | | |
| | Year | 1.795 | 0.598 | 3 | 4110.472 | 5.2 | 0.001 |
| | Bioclimatic Zone | 3.428 | 1.714 | 2 | 1155.731 | 14.891 | < 0.001 |
| | Year:Bioclimatic Zone | 2.58 | 0.43 | 6 | 4110.662 | 3.736 | 0.001 |
| Change in Functional Diversity (ΔFD) | | | | | | | |
| | Year Comparison | 1.979 | 0.99 | 2 | 2352 | 4.868 | 0.008 |
| | Bioclimatic Zone | 0.21 | 0.105 | 2 | 2352 | 0.516 | 0.597 |
| | Year Comparison:Bioclimatic Zone | 2.84 | 0.71 | 4 | 2352 | 3.493 | 0.008 |
| Resident species contribution to ΔFD (Res.) | | | | | | | |
| | Year Comparison | 3.282 | 1.641 | 2 | 2358 | 9.554 | < 0.001 |
| Turnover contribution to ΔFD | | | | | 2166 | | |
| | Year Comparison | 0.096 | 0.048 | 2 | 2166 | 3.558 | 0.029 |
| | Bioclimatic Zone | 0.072 | 0.036 | 2 | 2166 | 2.668 | 0.07 |
| | Year Comparison:Bioclimatic Zone | 0.162 | 0.04 | 4 | 2166 | 2.99 | 0.018 |
| Species Loss contribution to ΔFD (Loss) | | | | | | | |
| | Year Comparison | 0.11 | 0.055 | 2 | 1274.264 | 6.595 | 0.001 |
| | Bioclimatic Zone | 0.874 | 0.437 | 2 | 1003.03 | 52.59 | < 0.001 |
| | Year Comparison:Bioclimatic Zone | 0.091 | 0.023 | 4 | 1199.468 | 2.726 | 0.028 |
| Species Gain contribution to ΔFD (Gain) | | | | | | | |
| | Year Comparison | 0.287 | 0.144 | 2 | 1230.626 | 11.115 | < 0.001 |
| | Bioclimatic Zone | 0.864 | 0.432 | 2 | 689.135 | 33.416 | < 0.001 |
| Richness Loss contribution to ΔFD (Rich_L) | | | | | | | |
| | Year Comparison | 2.767 | 1.383 | 2 | 1515.848 | 35.18 | < 0.001 |
| | Bioclimatic Zone | 2.782 | 1.391 | 2 | 742.701 | 35.377 | < 0.001 |
| Richness Gain contribution to ΔFD (Rich_G) | | | | | | | |
| | Year Comparison | 0.641 | 0.32 | 2 | 1503.754 | 7.297 | 0.001 |
| | Bioclimatic Zone | 1.699 | 0.849 | 2 | 695.052 | 19.337 | < 0.001 |
| Compositional Loss contribution to ΔFD (Comp_L) | | | | | | | |
| | Year Comparison | 0.278 | 0.139 | 2 | 1288.619 | 4.847 | 0.008 |
| Compositional Gain contribution to ΔFD (Comp_G) | | | | | | | |
| | Year Comparison | 0.002 | 0.001 | 2 | 1539.186 | 0.034 | 0.966 |
| | Bioclimatic Zone | 0.084 | 0.042 | 2 | 671.853 | 1.624 | 0.198 |

Notes: Sum sq: Sum of Squares, Mean Sq: mean squared error, Num.DF: numerator degrees of freedom, Den.DF: denominator degrees of freedom.

**Table S2.** Post-hoc test results for models of Species Richness (SR), Functional Diversity (FD), Changes in FD (ΔFD), Turnover species' contribution to ΔFD, and Resident species' contribution to ΔFD (Res.) (Fig. 2, 3 & S1).

| Response | Contrast | Estimate | SE | DF | t-ratio | P-value |
|---|---|---|---|---|---|---|
| Species Richness (SR) | | | | | | |
| | South Boreal Zone | | | | | |
| | Year1985 - Year1995 | 0.172 | 0.139 | 1081.125 | 1.235 | 0.605 |
| | Year1985 - Year2006 | -0.166 | 0.235 | 1181.514 | -0.709 | 0.894 |
| | Year1985 - Year2023 | 1.143 | 0.137 | 1060.973 | 8.354 | < 0.001 |
| | Year1995 - Year2006 | -0.338 | 0.229 | 1081.125 | -1.478 | 0.451 |
| | Year1995 - Year2023 | 0.971 | 0.127 | 1060.973 | 7.669 | < 0.001 |
| | Year2006 - Year2023 | 1.31 | 0.227 | 1060.973 | 5.758 | < 0.001 |
| | Middle Boreal Zone | | | | | |
| | Year1985 - Year1995 | -0.044 | 0.129 | 1221.846 | -0.339 | 0.987 |
| | Year1985 - Year2006 | -0.22 | 0.222 | 1266.505 | -0.991 | 0.755 |
| | Year1985 - Year2023 | 0.124 | 0.13 | 1230.702 | 0.954 | 0.776 |
| | Year1995 - Year2006 | -0.176 | 0.22 | 1221.846 | -0.8 | 0.855 |
| | Year1995 - Year2023 | 0.167 | 0.127 | 1221.846 | 1.317 | 0.552 |
| | Year2006 - Year2023 | 0.343 | 0.221 | 1230.702 | 1.556 | 0.405 |
| | North Boreal Zone | | | | | |
| | Year1985 - Year1995 | -0.095 | 0.21 | 1459.875 | -0.454 | 0.969 |
| | Year1985 - Year2006 | -0.164 | 0.333 | 1498.226 | -0.492 | 0.961 |
| | Year1985 - Year2023 | -0.032 | 0.207 | 1253.193 | -0.155 | 0.999 |
| | Year1995 - Year2006 | -0.068 | 0.333 | 1459.875 | -0.206 | 0.997 |
| | Year1995 - Year2023 | 0.063 | 0.205 | 1253.193 | 0.308 | 0.99 |
| | Year2006 - Year2023 | 0.131 | 0.331 | 1253.193 | 0.397 | 0.979 |
| | 1985 | | | | | |
| | South Boreal Zone - Middle Boreal Zone | 2.308 | 0.283 | 1181.514 | 8.15 | < 0.001 |
| | South Boreal Zone - North Boreal Zone | 2.85 | 0.362 | 1181.514 | 7.87 | < 0.001 |
| | Middle Boreal Zone - North Boreal Zone | 0.542 | 0.351 | 1266.505 | 1.545 | 0.27 |
| | 1995 | | | | | |
| | South Boreal Zone - Middle Boreal Zone | 2.092 | 0.277 | 1081.125 | 7.561 | < 0.001 |
| | South Boreal Zone - North Boreal Zone | 2.583 | 0.359 | 1081.125 | 7.195 | < 0.001 |
| | Middle Boreal Zone - North Boreal Zone | 0.491 | 0.352 | 1221.846 | 1.394 | 0.344 |
| | 2006 | | | | | |
| | South Boreal Zone - Middle Boreal Zone | 2.254 | 0.385 | 2868.579 | 5.851 | < 0.001 |
| | South Boreal Zone - North Boreal Zone | 2.853 | 0.486 | 2868.579 | 5.874 | < 0.001 |
| | Middle Boreal Zone - North Boreal Zone | 0.598 | 0.477 | 3176.535 | 1.255 | 0.421 |
| | 2023 | | | | | |
| | South Boreal Zone - Middle Boreal Zone | 1.288 | 0.261 | 1060.973 | 4.941 | < 0.001 |
| | South Boreal Zone - North Boreal Zone | 1.675 | 0.336 | 1060.973 | 4.979 | < 0.001 |
| | Middle Boreal Zone - North Boreal Zone | 0.386 | 0.338 | 1230.702 | 1.143 | 0.488 |
| Functional Diversity (FD) | | | | | | |
| | South Boreal Zone | | | | | |
| | Year1985 - Year1995 | -0.056 | 0.017 | 3909.866 | -3.326 | 0.005 |
| | Year1985 - Year2006 | -0.068 | 0.027 | 4455.748 | -2.537 | 0.055 |
| | Year1985 - Year2023 | -0.059 | 0.017 | 4060.694 | -3.496 | 0.003 |
| | Year1995 - Year2006 | -0.013 | 0.026 | 4405.248 | -0.491 | 0.961 |
| | Year1995 - Year2023 | -0.003 | 0.016 | 3915.057 | -0.189 | 0.998 |

| | | | | | | |
|---|---|---|---|---|---|---|
| | Year2006 - Year2023 | 0.01 | 0.026 | 4445.928 | 0.378 | 0.982 |
| Middle Boreal Zone | | | | | | |
| | Year1985 - Year1995 | -0.043 | 0.02 | 3826.519 | -2.218 | 0.118 |
| | Year1985 - Year2006 | 0.044 | 0.032 | 4392.367 | 1.356 | 0.527 |
| | Year1985 - Year2023 | -0.08 | 0.02 | 3930.245 | -4.062 | < 0.001 |
| | Year1995 - Year2006 | 0.087 | 0.032 | 4379.698 | 2.719 | 0.033 |
| | Year1995 - Year2023 | -0.037 | 0.019 | 3873.806 | -1.904 | 0.226 |
| | Year2006 - Year2023 | -0.124 | 0.032 | 4408.266 | -3.855 | 0.001 |
| North Boreal Zone | | | | | | |
| | Year1985 - Year1995 | -0.041 | 0.034 | 3762.583 | -1.19 | 0.633 |
| | Year1985 - Year2006 | 0.07 | 0.052 | 4267.815 | 1.334 | 0.541 |
| | Year1985 - Year2023 | 0.036 | 0.033 | 4067.779 | 1.101 | 0.689 |
| | Year1995 - Year2006 | 0.11 | 0.052 | 4283.112 | 2.117 | 0.148 |
| | Year1995 - Year2023 | 0.077 | 0.033 | 4011.008 | 2.367 | 0.084 |
| | Year2006 - Year2023 | -0.033 | 0.051 | 4441.491 | -0.65 | 0.915 |
| 1985 | | | | | | |
| | South Boreal Zone - Middle Boreal Zone | 0.041 | 0.023 | 2451.823 | 1.788 | 0.174 |
| | South Boreal Zone - North Boreal Zone | 0.06 | 0.033 | 2806.328 | 1.826 | 0.161 |
| | Middle Boreal Zone - North Boreal Zone | 0.02 | 0.034 | 2826.475 | 0.58 | 0.831 |
| 1995 | | | | | | |
| | South Boreal Zone - Middle Boreal Zone | 0.053 | 0.022 | 2224.994 | 2.422 | 0.041 |
| | South Boreal Zone - North Boreal Zone | 0.075 | 0.032 | 2653.447 | 2.337 | 0.051 |
| | Middle Boreal Zone - North Boreal Zone | 0.023 | 0.034 | 2710.591 | 0.674 | 0.779 |
| 2006 | | | | | | |
| | South Boreal Zone - Middle Boreal Zone | 0.153 | 0.04 | 5170.395 | 3.838 | < 0.001 |
| | South Boreal Zone - North Boreal Zone | 0.199 | 0.056 | 5180.024 | 3.579 | 0.001 |
| | Middle Boreal Zone - North Boreal Zone | 0.046 | 0.058 | 5194.531 | 0.789 | 0.71 |
| 2023 | | | | | | |
| | South Boreal Zone - Middle Boreal Zone | 0.019 | 0.022 | 2220.897 | 0.868 | 0.661 |
| | South Boreal Zone - North Boreal Zone | 0.155 | 0.03 | 2271.939 | 5.211 | < 0.001 |
| | Middle Boreal Zone - North Boreal Zone | 0.136 | 0.031 | 2378.942 | 4.356 | < 0.001 |
| Change in Functional Diversity (ΔFD) | | | | | | |
| South Boreal Zone | | | | | | |
| | (1985 - 1995) - (1995 - 2006) | 0.02 | 0.034 | 2352 | 0.572 | 0.835 |
| | (1985 - 1995) - (2006 - 2023) | 0.049 | 0.035 | 2352 | 1.412 | 0.335 |
| | (1995 - 2006) - (2006 - 2023) | 0.029 | 0.043 | 2352 | 0.689 | 0.77 |
| Middle Boreal Zone | | | | | | |
| | (1985 - 1995) - (1995 - 2006) | 0.136 | 0.041 | 2352 | 3.319 | 0.003 |
| | (1985 - 1995) - (2006 - 2023) | -0.081 | 0.042 | 2352 | -1.942 | 0.127 |
| | (1995 - 2006) - (2006 - 2023) | -0.217 | 0.052 | 2352 | -4.172 | < 0.001 |
| North Boreal Zone | | | | | | |
| | (1985 - 1995) - (1995 - 2006) | 0.119 | 0.067 | 2352 | 1.79 | 0.173 |
| | (1985 - 1995) - (2006 - 2023) | 0.011 | 0.067 | 2352 | 0.166 | 0.985 |
| | (1995 - 2006) - (2006 - 2023) | -0.108 | 0.083 | 2352 | -1.31 | 0.389 |
| 1985 - 1995 | | | | | | |
| | South Boreal Zone - Middle Boreal Zone | -0.007 | 0.025 | 2352 | -0.262 | 0.963 |
| | South Boreal Zone - North Boreal Zone | 0.01 | 0.037 | 2352 | 0.275 | 0.959 |
| | Middle Boreal Zone - North Boreal Zone | 0.017 | 0.038 | 2352 | 0.441 | 0.898 |
| 1995 - 2006 | | | | | | |
| | South Boreal Zone - Middle Boreal Zone | 0.11 | 0.047 | 2352 | 2.334 | 0.051 |

| | | | | | |
|---|---|---|---|---|---|
| South Boreal Zone - North Boreal Zone | 0.11 | 0.065 | 2352 | 1.681 | 0.213 |
| Middle Boreal Zone - North Boreal Zone | 0 | 0.069 | 2352 | 0.002 | 1 |
| 2006 - 2023 | | | | | |
| South Boreal Zone - Middle Boreal Zone | -0.137 | 0.048 | 2352 | -2.841 | 0.013 |
| South Boreal Zone - North Boreal Zone | -0.028 | 0.066 | 2352 | -0.419 | 0.908 |
| Middle Boreal Zone - North Boreal Zone | 0.109 | 0.07 | 2352 | 1.569 | 0.259 |
| Resident species contribution to ΔFD (Res.) | | | | | |
| (1985 - 1995) - (1995 - 2006) | 0.087 | 0.022 | 2358 | 3.854 | < 0.001 |
| (1985 - 1995) - (2006 - 2023) | -0.026 | 0.023 | 2358 | -1.145 | 0.486 |
| (1995 - 2006) - (2006 - 2023) | -0.113 | 0.028 | 2358 | -3.997 | < 0.001 |
| Turnover contribution to ΔFD | | | | | |
| South Boreal Zone | | | | | |
| (1985 - 1995) - (1995 - 2006) | -0.015 | 0.009 | 2166 | -1.667 | 0.218 |
| (1985 - 1995) - (2006 - 2023) | 0.031 | 0.009 | 2166 | 3.445 | 0.002 |
| (1995 - 2006) - (2006 - 2023) | 0.046 | 0.011 | 2166 | 4.131 | < 0.001 |
| Middle Boreal Zone | | | | | |
| (1985 - 1995) - (1995 - 2006) | 0 | 0.011 | 2166 | 0.023 | 1 |
| (1985 - 1995) - (2006 - 2023) | -0.009 | 0.011 | 2166 | -0.841 | 0.678 |
| (1995 - 2006) - (2006 - 2023) | -0.01 | 0.014 | 2166 | -0.691 | 0.769 |
| North Boreal Zone | | | | | |
| (1985 - 1995) - (1995 - 2006) | 0.004 | 0.02 | 2166 | 0.178 | 0.983 |
| (1985 - 1995) - (2006 - 2023) | 0.002 | 0.02 | 2166 | 0.084 | 0.996 |
| (1995 - 2006) - (2006 - 2023) | -0.002 | 0.025 | 2166 | -0.078 | 0.997 |
| 1985 - 1995 | | | | | |
| South Boreal Zone - Middle Boreal Zone | -0.006 | 0.007 | 2166 | -0.892 | 0.645 |
| South Boreal Zone - North Boreal Zone | -0.012 | 0.011 | 2166 | -1.146 | 0.486 |
| Middle Boreal Zone - North Boreal Zone | -0.006 | 0.011 | 2166 | -0.571 | 0.836 |
| 1995 - 2006 | | | | | |
| South Boreal Zone - Middle Boreal Zone | 0.009 | 0.013 | 2166 | 0.735 | 0.743 |
| South Boreal Zone - North Boreal Zone | 0.006 | 0.019 | 2166 | 0.324 | 0.944 |
| Middle Boreal Zone - North Boreal Zone | -0.003 | 0.02 | 2166 | -0.146 | 0.988 |
| 2006 - 2023 | | | | | |
| South Boreal Zone - Middle Boreal Zone | -0.047 | 0.013 | 2166 | -3.66 | 0.001 |
| South Boreal Zone - North Boreal Zone | -0.042 | 0.019 | 2166 | -2.193 | 0.073 |
| Middle Boreal Zone - North Boreal Zone | 0.005 | 0.02 | 2166 | 0.238 | 0.969 |
| Species Loss contribution to ΔFD (Loss) | | | | | |
| South Boreal Zone | | | | | |
| (1985 - 1995) - (1995 - 2006) | 0.000 | 0.008 | 1386.929 | -0.047 | 0.999 |
| (1985 - 1995) - (2006 - 2023) | 0.057 | 0.008 | 1396.613 | 7.193 | < 0.001 |
| (1995 - 2006) - (2006 - 2023) | 0.057 | 0.009 | 751.311 | 6.109 | < 0.001 |
| Middle Boreal Zone | | | | | |
| (1985 - 1995) - (1995 - 2006) | -0.009 | 0.01 | 1413.751 | -0.847 | 0.674 |
| (1985 - 1995) - (2006 - 2023) | 0.027 | 0.01 | 1444.913 | 2.617 | 0.024 |
| (1995 - 2006) - (2006 - 2023) | 0.035 | 0.012 | 778.378 | 2.93 | 0.01 |
| North Boreal Zone | | | | | |
| (1985 - 1995) - (1995 - 2006) | 0.001 | 0.021 | 1514.413 | 0.029 | 1 |
| (1985 - 1995) - (2006 - 2023) | -0.002 | 0.021 | 1601.811 | -0.074 | 0.997 |
| (1995 - 2006) - (2006 - 2023) | -0.002 | 0.025 | 1037.638 | -0.085 | 0.996 |
| 1985 - 1995 | | | | | |

| | | | | | |
|---|---|---|---|---|---|
| | South Boreal Zone - Middle Boreal Zone | -0.037 | 0.007 | 922.423 | -5.272 | < 0.001 |
| | South Boreal Zone - North Boreal Zone | -0.085 | 0.012 | 1195.391 | -7.351 | < 0.001 |
| | Middle Boreal Zone - North Boreal Zone | -0.048 | 0.012 | 1215.875 | -4.048 | < 0.001 |
| 1995 - 2006 | | | | | | |
| | South Boreal Zone - Middle Boreal Zone | -0.045 | 0.012 | 1807.275 | -3.704 | 0.001 |
| | South Boreal Zone - North Boreal Zone | -0.084 | 0.021 | 1789.042 | -3.967 | < 0.001 |
| | Middle Boreal Zone - North Boreal Zone | -0.039 | 0.022 | 1790.547 | -1.782 | 0.176 |
| 2006 - 2023 | | | | | | |
| | South Boreal Zone - Middle Boreal Zone | -0.067 | 0.012 | 1816.288 | -5.549 | < 0.001 |
| | South Boreal Zone - North Boreal Zone | -0.144 | 0.02 | 1819.189 | -7.093 | < 0.001 |
| | Middle Boreal Zone - North Boreal Zone | -0.077 | 0.021 | 1818.83 | -3.63 | 0.001 |
| Species Gain contribution to ΔFD (Gain) | | | | | | |
| | (1985 - 1995) - (1995 - 2006) | 0.004 | 0.007 | 1511.959 | 0.533 | 0.855 |
| | (1985 - 1995) - (2006 - 2023) | -0.032 | 0.007 | 1506.7 | -4.377 | < 0.001 |
| | (1995 - 2006) - (2006 - 2023) | -0.036 | 0.009 | 901.395 | -4.065 | < 0.001 |
| | South Boreal Zone - Middle Boreal Zone | 0.032 | 0.007 | 620.074 | 4.605 | < 0.001 |
| | South Boreal Zone - North Boreal Zone | 0.084 | 0.011 | 807.346 | 7.691 | < 0.001 |
| | Middle Boreal Zone - North Boreal Zone | 0.052 | 0.011 | 823.087 | 4.622 | < 0.001 |
| Richness Loss contribution to ΔFD (Rich_L) | | | | | | |
| | (1985 - 1995) - (1995 - 2006) | -0.011 | 0.013 | 1613.444 | -0.852 | 0.671 |
| | (1985 - 1995) - (2006 - 2023) | 0.098 | 0.013 | 1610.274 | 7.841 | < 0.001 |
| | (1995 - 2006) - (2006 - 2023) | 0.109 | 0.015 | 1338.205 | 7.087 | < 0.001 |
| | South Boreal Zone - Middle Boreal Zone | -0.066 | 0.012 | 629.975 | -5.473 | < 0.001 |
| | South Boreal Zone - North Boreal Zone | -0.149 | 0.02 | 853.547 | -7.523 | < 0.001 |
| | Middle Boreal Zone - North Boreal Zone | -0.083 | 0.021 | 865.745 | -4.052 | < 0.001 |
| Richness Gain contribution to ΔFD (Rich_G) | | | | | | |
| | (1985 - 1995) - (1995 - 2006) | -0.003 | 0.013 | 1606.572 | -0.235 | 0.97 |
| | (1985 - 1995) - (2006 - 2023) | -0.05 | 0.013 | 1589.929 | -3.746 | 0.001 |
| | (1995 - 2006) - (2006 - 2023) | -0.047 | 0.016 | 1358.642 | -2.898 | 0.011 |
| | South Boreal Zone - Middle Boreal Zone | 0.026 | 0.012 | 620.048 | 2.115 | 0.088 |
| | South Boreal Zone - North Boreal Zone | 0.123 | 0.02 | 807.142 | 6.19 | < 0.001 |
| | Middle Boreal Zone - North Boreal Zone | 0.097 | 0.021 | 822.944 | 4.685 | < 0.001 |
| Compositional Loss contribution to ΔFD (Comp_L) | | | | | | |
| | (1985 - 1995) - (1995 - 2006) | 0.01 | 0.011 | 1560.59 | 0.894 | 0.644 |
| | (1985 - 1995) - (2006 - 2023) | -0.028 | 0.011 | 1570.099 | -2.646 | 0.022 |
| | (1995 - 2006) - (2006 - 2023) | -0.038 | 0.013 | 904.788 | -2.898 | 0.011 |
| Compositional Gain contribution to ΔFD (Comp_G) | | | | | | |
| | (1985 - 1995) - (1995 - 2006) | 0.001 | 0.01 | 1642.827 | 0.092 | 0.995 |
| | (1985 - 1995) - (2006 - 2023) | -0.002 | 0.01 | 1629.592 | -0.215 | 0.975 |
| | (1995 - 2006) - (2006 - 2023) | -0.003 | 0.012 | 1396.74 | -0.25 | 0.966 |
| | South Boreal Zone - Middle Boreal Zone | 0.014 | 0.009 | 588.81 | 1.598 | 0.247 |
| | South Boreal Zone - North Boreal Zone | -0.006 | 0.014 | 806.069 | -0.407 | 0.913 |
| | Middle Boreal Zone - North Boreal Zone | -0.02 | 0.015 | 822.03 | -1.349 | 0.369 |

Notes: SE: standard error, DF: degrees of freedom.

**Table S3.** Abbreviations and interpretation of the 5-parts Price components partitioning (Fig. 1, Eq. 2).

| Variable name | Notation | Description | Additional information |
|---|---|---|---|
| Functional Diversity | $FD, FD'$ | Rao's Q diversity measured using 9 traits at baseline / comparison time point | Sum of pairwise Euclidean distances between species, weighted by their relative abundance |
| Species number | $S, S'$ | Number of species at the baseline / comparison time point | Total number of species at a given time point |
| Common species number | $S_c$ | Number of species common to baseline and comparison time points | Species identity's overlap between consecutive time points for a same community (i.e., number of resident species) |
| Species-specific functional contribution | $Z_i\ , Z_i{}'$ | Contribution of species i to ΔFD at the baseline / comparison time point | Based on species dissimilarity with others in the community and their relative abundance (see Eq. 1) |
| Average species functional contribution | $\bar{Z}\ , \bar{Z}'$ | Average functional contribution per species at baseline / comparison time point | Sum of species-specific functional contribution divided by the number of species in the community at a given time point (resident and lost species at baseline, resident and gained species at comparison) |
| Average resident species functional contribution | $\bar{Z}_c\ , \bar{Z}'_c$ | Average functional contribution of the species common to baseline and comparison time points | Can differ between baseline and comparison time points, depending on the rest of the community changing around resident species (lost or gained species respectively) |
| Rich_L | $(S_c - S)\ \bar{Z}$ | Contribution of species loss to ΔFD, when species lost would be *average* species (or *random*), i.e., have a species-specific functional contribution equal to the average of the community | Equivalent to SRE_L in Fox and Kerr 2012, always negative. Isolates the effect of species loss on FD solely due to the number of species lost (also *random loss*). |
| Rich_G | $(S' - S_c)\ \bar{Z}'$ | Contribution of species gain to ΔFD, when species gained would be *average* species (or *random*), i.e., have a species-specific functional contribution equal to the average of the community | Equivalent to SRE_G in Fox and Kerr 2012, always positive. Isolates the effect of species gain on ΔFD solely due to the number of species gained (also *random gain*). |
| Comp_L | $S_c\ (\bar{Z}_c - \bar{Z})$ | Contribution of species loss to ΔFD, when considering only | Equivalent to SCE_L in Fox and Kerr 2012. Isolates the effect of species lost functional |

| | | | |
|---|---|---|---|
| | | the functional contribution of species lost | contribution on ΔFD, independently from their number (also *non-random loss*) |
| Comp_G | $-S_c\,(\bar{Z}'_c - \bar{Z}')$ | Contribution of species gain to ΔFD, when considering only the functional contribution species gained | Equivalent to SCE_G in Fox and Kerr 2012. Isolates the effect of species gained functional contribution on ΔFD, independently from their number (also *non-random gain*) |
| Res. | $S_c\,(\bar{Z}'_c - \bar{Z}_c)$ | Contribution to ΔFD of the differences in functional contribution of species in common between baseline and comparison | Equivalent to CDE in Fox and Kerr 2012. Captures the contribution of changes in the relative abundance and dissimilarity of resident species between baseline and comparison |

**Table S4.** Model results for species groups' (resident, lost and gained species) Dissimilarity, Mean Relative Abundance and Number over consecutive survey years (Year Comparison) and Bioclimatic Zones (Fig. 4).

| Response | Predictor | Sum Sq. | Mean Sq. | Num.DF | Den.DF | F-value | P-value |
|---|---|---|---|---|---|---|---|
| Dissimilarity among resident sp. | | | | | | | |
| | Bioclimatic Zone | 4.499 | 2.25 | 2 | 755.36 | 17.639 | < 0.001 |
| Dissimilarity among lost sp. | | | | | | | |
| | Bioclimatic Zone | 1.395 | 0.697 | 2 | 875.14 | 9.535 | < 0.001 |
| Dissimilarity among gained sp. | | | | | | | |
| | Bioclimatic Zone | 0.189 | 0.094 | 2 | 890.792 | 1.436 | 0.238 |
| Dissimilarity between gained and resident sp. | | | | | | | |
| | Bioclimatic Zone | 1.169 | 0.585 | 2 | 679.916 | 32.544 | < 0.001 |
| Dissimilarity between lost and resident sp. | | | | | | | |
| | Year Comparison | 0.09 | 0.045 | 2 | 1002.334 | 2.409 | 0.09 |
| | Bioclimatic Zone | 1.757 | 0.878 | 2 | 710.653 | 47.159 | < 0.001 |
| Relative Abundance of resident sp. | | | | | | | |
| | Year Comparison | 0.544 | 0.272 | 2 | 917.912 | 5.863 | 0.003 |
| | Bioclimatic Zone | 3.012 | 1.506 | 2 | 985.944 | 32.447 | < 0.001 |
| | Year Comparison:Bioclimatic Zone | 0.378 | 0.095 | 4 | 922.864 | 2.038 | 0.087 |
| Relative Abundance of lost sp. | | | | | | | |
| | Year Comparison | 2.564 | 1.282 | 2 | 1264.454 | 0.964 | 0.382 |
| | Bioclimatic Zone | 92.086 | 46.043 | 2 | 971.459 | 34.624 | < 0.001 |
| | Year Comparison:Bioclimatic Zone | 21.524 | 5.381 | 4 | 1186.938 | 4.047 | 0.003 |
| Relative Abundance of gained sp. | | | | | | | |
| | Year Comparison | 29.894 | 14.947 | 2 | 1314.505 | 10.013 | < 0.001 |
| | Bioclimatic Zone | 51.422 | 25.711 | 2 | 850.182 | 17.224 | < 0.001 |
| | Year Comparison:Bioclimatic Zone | 11.883 | 2.971 | 4 | 1257.663 | 1.99 | 0.094 |

| Response | Predictor | Nb.par. | logLik. | DF | Deviance | Chi-sq. | P-value |
|---|---|---|---|---|---|---|---|
| Number of resident sp. | | | | | | | |
| | Year Comparison | 7 | -5840.2 | 2 | 11680 | 49.892 | 1.47E-11 |
| | Bioclimatic Zone | 7 | -5840.2 | 2 | 11680 | 63.27 | 1.83E-14 |
| Number of lost sp. | | | | | | | |
| | Year Comparison | 7 | -4831.6 | 2 | 9663.3 | 153.34 | < 2.2e-16 |
| | Bioclimatic Zone | 7 | -4831.6 | 2 | 9663.3 | 207.74 | < 2.2e-16 |
| | Year Comparison:Bioclimatic Zone | 11 | -4826.6 | 4 | 9653.3 | 10.027 | 0.03997 |
| Number of gained sp. | | | | | | | |
| | Year Comparison | 7 | -4924 | 2 | 9847.9 | 5.5387 | 0.0627 |
| | Bioclimatic Zone | 7 | -4924 | 2 | 9847.9 | 126.11 | < 2.2e-16 |

Notes: Sum sq: Sum of Squares, Mean Sq: mean squared error, Num.DF: numerator degrees of freedom, Den.DF: denominator degrees of freedom, Nb.par: number of parameters, logLik.: log-likelihood.

**Table S5.** Post-hoc test results for models of species groups' (resident, lost and gained species) Dissimilarity, Relative Abundance and Number (Fig. 4).

| Response | Contrast | Estimate | SE | DF | t-ratio | P-value |
|---|---|---|---|---|---|---|
| Dissimilarity within Resident sp. | | | | | | |
| | South Boreal Zone - Middle Boreal Zone | -0.216 | 0.037 | 722.056 | -5.83 | < 0.001 |
| | South Boreal Zone - North Boreal Zone | -0.036 | 0.054 | 748.428 | -0.669 | 0.781 |
| | Middle Boreal Zone - North Boreal Zone | 0.18 | 0.055 | 768.981 | 3.243 | 0.004 |
| Dissimilarity within Lost sp. | | | | | | |
| | South Boreal Zone - Middle Boreal Zone | -0.289 | 0.068 | 758.685 | -4.244 | < 0.001 |
| | South Boreal Zone - North Boreal Zone | -0.196 | 0.136 | 758.685 | -1.449 | 0.316 |
| | Middle Boreal Zone - North Boreal Zone | 0.093 | 0.142 | 877.353 | 0.654 | 0.79 |
| Dissimilarity within Gained sp. | | | | | | |
| | South Boreal Zone - Middle Boreal Zone | -0.097 | 0.067 | 799.823 | -1.455 | 0.313 |
| | South Boreal Zone - North Boreal Zone | -0.147 | 0.133 | 799.823 | -1.102 | 0.513 |
| | Middle Boreal Zone - North Boreal Zone | -0.05 | 0.138 | 839.287 | -0.363 | 0.93 |
| Dissimilarity between Lost-Res | | | | | | |
| | South Boreal Zone | | | | | |
| | (1985 - 1995) - (1995 - 2006) | -0.055 | 0.041 | 750.076 | -1.354 | 0.366 |
| | (1985 - 1995) - (2006 - 2023) | -0.083 | 0.041 | 750.076 | -2.058 | 0.099 |
| | (1995 - 2006) - (2006 - 2023) | -0.028 | 0.047 | 1666.387 | -0.591 | 0.825 |
| | Middle Boreal Zone | | | | | |
| | (1985 - 1995) - (1995 - 2006) | -0.061 | 0.045 | 777.708 | -1.353 | 0.366 |
| | (1985 - 1995) - (2006 - 2023) | -0.091 | 0.044 | 777.708 | -2.057 | 0.1 |
| | (1995 - 2006) - (2006 - 2023) | -0.031 | 0.052 | 1526.858 | -0.591 | 0.825 |
| | North Boreal Zone | | | | | |
| | (1985 - 1995) - (1995 - 2006) | -0.06 | 0.044 | 949.114 | -1.353 | 0.366 |
| | (1985 - 1995) - (2006 - 2023) | -0.09 | 0.044 | 949.114 | -2.057 | 0.1 |
| | (1995 - 2006) - (2006 - 2023) | -0.03 | 0.051 | 1225.407 | -0.591 | 0.825 |
| | 1985 - 1995 | | | | | |
| | South Boreal Zone - Middle Boreal Zone | -0.411 | 0.044 | 750.076 | -9.253 | < 0.001 |
| | South Boreal Zone - North Boreal Zone | -0.343 | 0.073 | 750.076 | -4.685 | < 0.001 |
| | Middle Boreal Zone - North Boreal Zone | 0.068 | 0.077 | 777.708 | 0.884 | 0.651 |
| | 1995 - 2006 | | | | | |
| | South Boreal Zone - Middle Boreal Zone | -0.416 | 0.045 | 1542.014 | -9.225 | < 0.001 |
| | South Boreal Zone - North Boreal Zone | -0.347 | 0.074 | 1242.514 | -4.682 | < 0.001 |
| | Middle Boreal Zone - North Boreal Zone | 0.069 | 0.078 | 1242.514 | 0.884 | 0.651 |
| | 2006 - 2023 | | | | | |
| | South Boreal Zone - Middle Boreal Zone | -0.418 | 0.045 | 1526.858 | -9.22 | < 0.001 |
| | South Boreal Zone - North Boreal Zone | -0.349 | 0.075 | 1225.407 | -4.683 | < 0.001 |
| | Middle Boreal Zone - North Boreal Zone | 0.069 | 0.078 | 1225.407 | 0.884 | 0.651 |
| Dissimilarity between Gained-Res | | | | | | |
| | South Boreal Zone - Middle Boreal Zone | -0.326 | 0.042 | 595.493 | -7.692 | < 0.001 |
| | South Boreal Zone - North Boreal Zone | -0.272 | 0.067 | 595.493 | -4.038 | < 0.001 |
| | Middle Boreal Zone - North Boreal Zone | 0.054 | 0.071 | 668.128 | 0.768 | 0.723 |
| Abundance of Resident sp. | | | | | | |
| | South Boreal Zone | | | | | |
| | (1985 - 1995) - (1995 - 2006) | 0.001 | 0.002 | 766.89 | 0.36 | 0.931 |
| | (1985 - 1995) - (2006 - 2023) | -0.011 | 0.003 | 766.89 | -4.109 | < 0.001 |
| | (1995 - 2006) - (2006 - 2023) | -0.011 | 0.002 | 1595.722 | -4.62 | < 0.001 |

| | | | | | |
|---|---|---|---|---|---|
| Middle Boreal Zone | | | | | |
| (1985 - 1995) - (1995 - 2006) | -0.004 | 0.004 | 791.769 | -1.191 | 0.459 |
| (1985 - 1995) - (2006 - 2023) | -0.007 | 0.004 | 791.769 | -1.938 | 0.129 |
| (1995 - 2006) - (2006 - 2023) | -0.003 | 0.004 | 1629.109 | -0.783 | 0.714 |
| North Boreal Zone | | | | | |
| (1985 - 1995) - (1995 - 2006) | -0.006 | 0.006 | 840.043 | -0.945 | 0.612 |
| (1985 - 1995) - (2006 - 2023) | -0.005 | 0.006 | 840.043 | -0.837 | 0.68 |
| (1995 - 2006) - (2006 - 2023) | 0.001 | 0.006 | 1573.503 | 0.1 | 0.994 |
| 1985 - 1995 | | | | | |
| South Boreal Zone - Middle Boreal Zone | -0.029 | 0.004 | 766.89 | -7.232 | < 0.001 |
| South Boreal Zone - North Boreal Zone | -0.037 | 0.006 | 766.89 | -5.736 | < 0.001 |
| Middle Boreal Zone - North Boreal Zone | -0.008 | 0.007 | 791.769 | -1.112 | 0.507 |
| 1995 - 2006 | | | | | |
| South Boreal Zone - Middle Boreal Zone | -0.034 | 0.005 | 1595.722 | -6.351 | < 0.001 |
| South Boreal Zone - North Boreal Zone | -0.043 | 0.008 | 1573.503 | -5.145 | < 0.001 |
| Middle Boreal Zone - North Boreal Zone | -0.009 | 0.009 | 1573.503 | -0.985 | 0.586 |
| 2006 - 2023 | | | | | |
| South Boreal Zone - Middle Boreal Zone | -0.026 | 0.006 | 1605.33 | -4.504 | < 0.001 |
| South Boreal Zone - North Boreal Zone | -0.031 | 0.008 | 1579.047 | -3.644 | 0.001 |
| Middle Boreal Zone - North Boreal Zone | -0.005 | 0.009 | 1579.047 | -0.584 | 0.829 |
| Abundance of Lost sp. | | | | | |
| South Boreal Zone | | | | | |
| (1985 - 1995) - (1995 - 2006) | -0.002 | 0.001 | 825.056 | -1.29 | 0.401 |
| (1985 - 1995) - (2006 - 2023) | -0.006 | 0.002 | 825.056 | -4.05 | < 0.001 |
| (1995 - 2006) - (2006 - 2023) | -0.005 | 0.002 | 1811.005 | -2.756 | 0.016 |
| Middle Boreal Zone | | | | | |
| (1985 - 1995) - (1995 - 2006) | 0.001 | 0.001 | 945.212 | 0.894 | 0.644 |
| (1985 - 1995) - (2006 - 2023) | -0.001 | 0.001 | 945.212 | -0.772 | 0.72 |
| (1995 - 2006) - (2006 - 2023) | -0.002 | 0.001 | 1812.519 | -1.382 | 0.351 |
| North Boreal Zone | | | | | |
| (1985 - 1995) - (1995 - 2006) | -0.002 | 0.002 | 1271.513 | -1.273 | 0.411 |
| (1985 - 1995) - (2006 - 2023) | 0.001 | 0.001 | 1271.513 | 1.43 | 0.326 |
| (1995 - 2006) - (2006 - 2023) | 0.004 | 0.002 | 1800.833 | 2.015 | 0.109 |
| 1985 - 1995 | | | | | |
| South Boreal Zone - Middle Boreal Zone | 0.003 | 0.001 | 825.056 | 3.692 | 0.001 |
| South Boreal Zone - North Boreal Zone | 0.006 | 0.001 | 825.056 | 7.013 | < 0.001 |
| Middle Boreal Zone - North Boreal Zone | 0.003 | 0.001 | 945.212 | 3.845 | < 0.001 |
| 1995 - 2006 | | | | | |
| South Boreal Zone - Middle Boreal Zone | 0.005 | 0.001 | 1811.005 | 3.673 | 0.001 |
| South Boreal Zone - North Boreal Zone | 0.005 | 0.002 | 1800.833 | 2.469 | 0.036 |
| Middle Boreal Zone - North Boreal Zone | 0 | 0.002 | 1800.833 | -0.022 | 1 |
| 2006 - 2023 | | | | | |
| South Boreal Zone - Middle Boreal Zone | 0.008 | 0.002 | 1818.524 | 4.493 | < 0.001 |
| South Boreal Zone - North Boreal Zone | 0.014 | 0.002 | 1822.118 | 7.864 | < 0.001 |
| Middle Boreal Zone - North Boreal Zone | 0.005 | 0.001 | 1818.524 | 4.078 | < 0.001 |
| Abundance of Gained sp. | | | | | |
| South Boreal Zone | | | | | |
| (1985 - 1995) - (1995 - 2006) | -0.001 | 0.001 | 884.38 | -0.532 | 0.855 |
| (1985 - 1995) - (2006 - 2023) | -0.006 | 0.002 | 884.38 | -3.493 | 0.001 |

| | | | | | |
|---|---|---|---|---|---|
| (1995 - 2006) - (2006 - 2023) | -0.006 | 0.002 | 1815.232 | -2.836 | 0.013 |
| Middle Boreal Zone | | | | | |
| (1985 - 1995) - (1995 - 2006) | 0.003 | 0.001 | 998.595 | 2.557 | 0.029 |
| (1985 - 1995) - (2006 - 2023) | -0.004 | 0.002 | 998.595 | -2.49 | 0.035 |
| (1995 - 2006) - (2006 - 2023) | -0.007 | 0.002 | 1813.253 | -3.866 | < 0.001 |
| North Boreal Zone | | | | | |
| (1985 - 1995) - (1995 - 2006) | 0.003 | 0.001 | 1314.825 | 2.15 | 0.08 |
| (1985 - 1995) - (2006 - 2023) | -0.001 | 0.002 | 1314.825 | -0.389 | 0.92 |
| (1995 - 2006) - (2006 - 2023) | -0.004 | 0.002 | 1810.284 | -1.798 | 0.17 |
| 1985 - 1995 | | | | | |
| South Boreal Zone - Middle Boreal Zone | 0.002 | 0.001 | 884.38 | 2.354 | 0.049 |
| South Boreal Zone - North Boreal Zone | 0.004 | 0.001 | 884.38 | 3.139 | 0.005 |
| Middle Boreal Zone - North Boreal Zone | 0.002 | 0.001 | 998.595 | 1.363 | 0.361 |
| 1995 - 2006 | | | | | |
| South Boreal Zone - Middle Boreal Zone | 0.006 | 0.001 | 1813.253 | 3.801 | < 0.001 |
| South Boreal Zone - North Boreal Zone | 0.008 | 0.002 | 1814.388 | 4.757 | < 0.001 |
| Middle Boreal Zone - North Boreal Zone | 0.002 | 0.001 | 1813.253 | 1.563 | 0.262 |
| 2006 - 2023 | | | | | |
| South Boreal Zone - Middle Boreal Zone | 0.004 | 0.002 | 1815.232 | 1.739 | 0.191 |
| South Boreal Zone - North Boreal Zone | 0.009 | 0.003 | 1810.284 | 3.492 | 0.001 |
| Middle Boreal Zone - North Boreal Zone | 0.005 | 0.003 | 1810.284 | 1.96 | 0.123 |
| Number of Resident sp. | | | | | |
| South Boreal Zone | | | | | |
| (1985 - 1995) - (1995 - 2006) | 0.279 | 0.176 | Inf | 1.585 | 0.2522 |
| (1985 - 1995) - (2006 - 2023) | 1.229 | 0.169 | Inf | 7.258 | <.0001 |
| (1995 - 2006) - (2006 - 2023) | 0.95 | 0.193 | Inf | 4.92 | <.0001 |
| Middle Boreal Zone | | | | | |
| (1985 - 1995) - (1995 - 2006) | 0.231 | 0.145 | Inf | 1.586 | 0.2517 |
| (1985 - 1995) - (2006 - 2023) | 1.017 | 0.14 | Inf | 7.259 | <.0001 |
| (1995 - 2006) - (2006 - 2023) | 0.786 | 0.16 | Inf | 4.912 | <.0001 |
| North Boreal Zone | | | | | |
| (1985 - 1995) - (1995 - 2006) | 0.222 | 0.141 | Inf | 1.583 | 0.2529 |
| (1985 - 1995) - (2006 - 2023) | 0.981 | 0.138 | Inf | 7.089 | <.0001 |
| (1995 - 2006) - (2006 - 2023) | 0.758 | 0.156 | Inf | 4.864 | <.0001 |
| 1985 - 1995 | | | | | |
| South Boreal Zone - Middle Boreal Zone | 1.438 | 0.2 | Inf | 7.206 | <.0001 |
| South Boreal Zone - North Boreal Zone | 1.683 | 0.273 | Inf | 6.174 | <.0001 |
| Middle Boreal Zone - North Boreal Zone | 0.245 | 0.272 | Inf | 0.902 | 0.6392 |
| 1995 - 2006 | | | | | |
| South Boreal Zone - Middle Boreal Zone | 1.39 | 0.194 | Inf | 7.174 | <.0001 |
| South Boreal Zone - North Boreal Zone | 1.627 | 0.265 | Inf | 6.145 | <.0001 |
| Middle Boreal Zone - North Boreal Zone | 0.237 | 0.263 | Inf | 0.902 | 0.6393 |
| 2006 - 2023 | | | | | |
| South Boreal Zone - Middle Boreal Zone | 1.226 | 0.171 | Inf | 7.165 | <.0001 |
| South Boreal Zone - North Boreal Zone | 1.435 | 0.234 | Inf | 6.138 | <.0001 |
| Middle Boreal Zone - North Boreal Zone | 0.209 | 0.232 | Inf | 0.901 | 0.6394 |
| Number of Lost sp. | | | | | |
| South Boreal Zone | | | | | |
| (1985 - 1995) - (1995 - 2006) | 0.287 | 0.131 | Inf | 2.187 | 0.0734 |

| | | | | | |
|---|---|---|---|---|---|
| (1985 - 1995) - (2006 - 2023) | -1.3583 | 0.179 | Inf | -7.569 | <.0001 |
| (1995 - 2006) - (2006 - 2023) | -1.6453 | 0.173 | Inf | -9.52 | <.0001 |
| Middle Boreal Zone | | | | | |
| (1985 - 1995) - (1995 - 2006) | -0.0478 | 0.123 | Inf | -0.388 | 0.9204 |
| (1985 - 1995) - (2006 - 2023) | -0.6842 | 0.151 | Inf | -4.524 | <.0001 |
| (1995 - 2006) - (2006 - 2023) | -0.6364 | 0.155 | Inf | -4.097 | 0.0001 |
| North Boreal Zone | | | | | |
| (1985 - 1995) - (1995 - 2006) | 0.3159 | 0.112 | Inf | 2.827 | 0.0131 |
| (1985 - 1995) - (2006 - 2023) | -0.1287 | 0.144 | Inf | -0.891 | 0.6462 |
| (1995 - 2006) - (2006 - 2023) | -0.4446 | 0.151 | Inf | -2.95 | 0.0089 |
| 1985 - 1995 | | | | | |
| South Boreal Zone - Middle Boreal Zone | 1.104 | 0.127 | Inf | 8.696 | <.0001 |
| South Boreal Zone - North Boreal Zone | 1.897 | 0.13 | Inf | 14.643 | <.0001 |
| Middle Boreal Zone - North Boreal Zone | 0.793 | 0.111 | Inf | 7.165 | <.0001 |
| 1995 - 2006 | | | | | |
| South Boreal Zone - Middle Boreal Zone | 0.769 | 0.185 | Inf | 4.16 | 0.0001 |
| South Boreal Zone - North Boreal Zone | 1.926 | 0.167 | Inf | 11.548 | <.0001 |
| Middle Boreal Zone - North Boreal Zone | 1.156 | 0.159 | Inf | 7.25 | <.0001 |
| 2006 - 2023 | | | | | |
| South Boreal Zone - Middle Boreal Zone | 1.778 | 0.259 | Inf | 6.856 | <.0001 |
| South Boreal Zone - North Boreal Zone | 3.127 | 0.249 | Inf | 12.56 | <.0001 |
| Middle Boreal Zone - North Boreal Zone | 1.348 | 0.217 | Inf | 6.218 | <.0001 |
| Number of Gained sp. | | | | | |
| South Boreal Zone | | | | | |
| (1985 - 1995) - (1995 - 2006) | -0.2387 | 0.108 | Inf | -2.214 | 0.0689 |
| (1985 - 1995) - (2006 - 2023) | -0.1644 | 0.107 | Inf | -1.53 | 0.2768 |
| (1995 - 2006) - (2006 - 2023) | 0.0743 | 0.114 | Inf | 0.655 | 0.7896 |
| Middle Boreal Zone | | | | | |
| (1985 - 1995) - (1995 - 2006) | -0.1606 | 0.0728 | Inf | -2.207 | 0.07 |
| (1985 - 1995) - (2006 - 2023) | -0.1106 | 0.0724 | Inf | -1.527 | 0.278 |
| (1995 - 2006) - (2006 - 2023) | 0.05 | 0.0764 | Inf | 0.655 | 0.7897 |
| North Boreal Zone | | | | | |
| (1985 - 1995) - (1995 - 2006) | -0.0848 | 0.0389 | Inf | -2.182 | 0.0743 |
| (1985 - 1995) - (2006 - 2023) | -0.0584 | 0.0384 | Inf | -1.52 | 0.2815 |
| (1995 - 2006) - (2006 - 2023) | 0.0264 | 0.0404 | Inf | 0.654 | 0.7901 |
| 1985 - 1995 | | | | | |
| South Boreal Zone - Middle Boreal Zone | 0.82 | 0.123 | Inf | 6.658 | <.0001 |
| South Boreal Zone - North Boreal Zone | 1.616 | 0.125 | Inf | 12.961 | <.0001 |
| Middle Boreal Zone - North Boreal Zone | 0.796 | 0.111 | Inf | 7.175 | <.0001 |
| 1995 - 2006 | | | | | |
| South Boreal Zone - Middle Boreal Zone | 0.898 | 0.137 | Inf | 6.544 | <.0001 |
| South Boreal Zone - North Boreal Zone | 1.77 | 0.147 | Inf | 12.079 | <.0001 |
| Middle Boreal Zone - North Boreal Zone | 0.872 | 0.125 | Inf | 6.995 | <.0001 |
| 2006 - 2023 | | | | | |
| South Boreal Zone - Middle Boreal Zone | 0.873 | 0.134 | Inf | 6.535 | <.0001 |
| South Boreal Zone - North Boreal Zone | 1.722 | 0.143 | Inf | 12.018 | <.0001 |
| Middle Boreal Zone - North Boreal Zone | 0.848 | 0.122 | Inf | 6.981 | <.0001 |

Notes: SE: standard error, DF: degrees of freedom, Inf: infinite degrees of freedom used for generalized linear model under the assumption of asymptotic normality.

**Table S6.** Model results for functional traits differences between species groups (resident, lost and gained species; Fig. S2).

| Response | Predictor | Sum Sq. | Mean Sq. | Num.DF | Den.DF | F-value | P-value |
|---|---|---|---|---|---|---|---|
| Leaf Dry Matter Content (LDMC) | species group | 39.363 | 19.681 | 2 | 4891.931 | 529.271 | < 0.001 |
| Leaf Carbon | species group | 935594.5 | 467797.3 | 2 | 4918.447 | 844.488 | < 0.001 |
| Leaf Nitrogen | species group | 5462.144 | 2731.072 | 2 | 5009.104 | 248.25 | < 0.001 |
| Leaf Phosphorus | species group | 184.254 | 92.127 | 2 | 5066.444 | 530.655 | < 0.001 |
| Plant height | species group | 371.622 | 185.811 | 2 | 4989.782 | 645.314 | < 0.001 |
| Rooting depth | species group | 619.081 | 309.541 | 2 | 5185.775 | 1015.981 | < 0.001 |
| Seed dry mass | species group | 114.254 | 57.127 | 2 | 5129.273 | 38.893 | < 0.001 |
| Specific Leaf Area (SLA) | species group | 31.573 | 15.787 | 2 | 4832.621 | 197.039 | < 0.001 |
| Specific Root Length (SRL) | species group | 129276.2 | 64638.09 | 2 | 5180.087 | 14.402 | < 0.001 |

Notes: Sum sq: Sum of Squares, Mean Sq: mean squared error, Num.DF: numerator degrees of freedom, Den.DF: denominator degrees of freedom.

**Table S7.** Post-hoc test results for models of functional traits differences between species groups (resident, lost and gained species; Fig. S2). See Table S6 for trait abbreviations.

| Response | Contrast | Estimate | SE | DF | t-ratio | P-value |
|---|---|---|---|---|---|---|
| LDMC | Gained - Lost | -0.008 | 0.002 | 2.014 | -4.299 | 0.089 |
| | Gained - Resident | -0.055 | 0.004 | 2.014 | -14.569 | 0.008 |
| | Lost - Resident | -0.047 | 0.003 | 2.014 | -13.92 | 0.009 |
| Leaf C | Gained - Lost | -2.718 | 0.78 | 4893.747 | -3.483 | 0.001 |
| | Gained - Resident | -27.015 | 0.739 | 4878.252 | -36.538 | < 0.001 |
| | Lost - Resident | -24.297 | 0.737 | 4869.228 | -32.988 | < 0.001 |
| Leaf N | Gained - Lost | 0.428 | 0.11 | 4991.016 | 3.901 | < 0.001 |
| | Gained - Resident | 2.146 | 0.104 | 4982.067 | 20.657 | < 0.001 |
| | Lost - Resident | 1.718 | 0.103 | 4972.263 | 16.6 | < 0.001 |
| Leaf P | Gained - Lost | 0.009 | 0.014 | 5036 | 0.665 | 0.784 |
| | Gained - Resident | 0.364 | 0.013 | 5029.362 | 27.947 | < 0.001 |
| | Lost - Resident | 0.355 | 0.013 | 5019.403 | 27.342 | < 0.001 |
| Plant height | Gained - Lost | 0.007 | 0.003 | 2.075 | 2.218 | 0.265 |
| | Gained - Resident | 0.079 | 0.007 | 2.044 | 11.751 | 0.012 |
| | Lost - Resident | 0.071 | 0.006 | 2.044 | 11.617 | 0.012 |
| Rooting depth | Gained - Lost | -0.008 | 0.005 | 2.262 | -1.731 | 0.362 |
| | Gained - Resident | 0.125 | 0.006 | 2.157 | 19.255 | 0.003 |
| | Lost - Resident | 0.133 | 0.007 | 2.157 | 19.522 | 0.003 |
| Seed dry mass | Gained - Lost | 0.015 | 0.026 | 2.215 | 0.589 | 0.838 |
| | Gained - Resident | 0.166 | 0.028 | 2.127 | 5.928 | 0.043 |
| | Lost - Resident | 0.151 | 0.027 | 2.127 | 5.643 | 0.048 |
| SLA | Gained - Lost | 1.089 | 0.208 | 2.009 | 5.241 | 0.062 |
| | Gained - Resident | 2.916 | 0.361 | 2.009 | 8.083 | 0.027 |
| | Lost - Resident | 1.827 | 0.252 | 2.009 | 7.25 | 0.033 |
| SRL | Gained - Lost | 10.541 | 2.21 | 5060.1 | 4.77 | < 0.001 |
| | Gained - Resident | 0.94 | 2.094 | 5055.019 | 0.449 | 0.895 |
| | Lost - Resident | -9.6 | 2.087 | 5044.959 | -4.601 | < 0.001 |

**Table S8.** Functional traits range and imputation errors. See Table S6 for trait abbreviations.

| Trait | Trait unit | min. | max. | mean | sd. | Percentage imputed | RMSE | Overall NRMSE |
|---|---|---|---|---|---|---|---|---|
| Plant height | meters | 0.004 | 3.160 | 0.429 | 0.427 | 2.683 | 0.374 | 0.396 |
| SLA | mm2/mg | 3.428 | 104.135 | 24.079 | 12.880 | 7.805 | 11.030 | |
| LDMC | g/g | 0.065 | 0.637 | 0.251 | 0.090 | 9.756 | 0.059 | |
| Seed dry mass | mg | 0.0001 | 70.380 | 2.352 | 5.796 | 12.195 | 5.239 | |
| Leaf C | mg/g | 265.000 | 555.355 | 434.422 | 33.536 | 20.976 | 28.972 | |
| Leaf N | mg/g | 7.860 | 53.751 | 22.737 | 7.656 | 15.122 | 6.500 | |
| Leaf P | mg/g | 0.272 | 7.000 | 2.255 | 0.988 | 25.854 | 0.930 | |
| Rooting depth | meters | 0.007 | 2.438 | 0.381 | 0.275 | 55.366 | 0.352 | |
| SRL | m/g | 4.576 | 1033.316 | 205.755 | 108.182 | 67.317 | 169.621 | |

Notes: sd: standard deviation; (N)RMSE: (normalised) root mean square error.
